# Epigenetic alterations at distal enhancers are linked to proliferation in human breast cancer

**DOI:** 10.1101/2021.04.14.439799

**Authors:** Jørgen Ankill, Miriam Ragle Aure, Sunniva Bjørklund, Severin Langberg, Oslo Breast Cancer Research Consortium (OSBREAC), Vessela N. Kristensen, Valeria Vitelli, Xavier Tekpli, Thomas Fleischer

## Abstract

Breast cancer is a highly heterogeneous disease driven by multiple factors including genetic and epigenetic alterations. DNA methylation patterns have been shown to be altered on a genome-wide scale and previous studies have highlighted the critical role of aberrant DNA methylation on gene expression and breast cancer pathogenesis. Here, we perform genome-wide expression-methylation Quantitative Trait Loci (emQTL), a method for integration of CpG methylation and gene expression to identify disease-driving genes under epigenetic control. By grouping these emQTLs by biclustering we identify associations representing important biological processes associated with breast cancer pathogenesis such as proliferation and tumor infiltrating fibroblasts. We report hypomethylation at enhancers carrying transcription factor binding sites of key proliferation-driving transcription factors such as CEBP-β, FOSL1, and FOSL2, with concomitant high expression of cell cycle- and proliferation-related genes in aggressive breast tumors. The identified CpGs and genes were found to be connected through chromatin loops, together indicating that proliferation in aggressive breast tumors is under epigenetic regulation by DNA methylation. Interestingly, there was a significant correlation between proliferation-related DNA methylation and gene expression also within subtypes of breast cancer, thereby showing that varying proliferation may be explained by epigenetic profiles across breast cancer subtypes. Indeed, the identified proliferation gene signature was prognostic both in the Luminal A and Luminal B subtypes. Taken together, we show that proliferation in breast cancer is linked to hypomethylation at specific enhancers and transcription factor binding mediated through chromatin loops.

## Introduction

Epigenetic alterations, such as DNA methylation, have recently emerged as a hallmark of many cancer types including breast cancer. Previous studies have shown that changes in DNA methylation patterns are present already in pre-invasive lesions thereby suggesting that such alterations occur early during breast cancer carcinogenesis^1-3^. DNA methylation has been predominantly reported to be implicated in gene repression through promoter methylation^4^, however, we have shown that DNA methylation at CpGs up to 100 kb away from gene transcription start sites could be associated with its expression^5^. Furthermore, a major portion of the aberrantly methylated DNA observed in breast cancers occurs in intergenic regions. Altogether, this suggests that DNA methylation at distal cis-regulatory regions such as enhancers may be an important contributor to breast cancer development and progression^5,6^.

Enhancers are *cis*-acting DNA sequences involved in transcriptional regulation. This process is mediated by binding of cell type specific transcription factors (TFs) and formation of physical interactions between enhancers and promoters of their associated genes^7,8^. TFs are key proteins involved in regulation of gene expression and are linked to different functions dependent on where they bind in the genome. While some TFs activate gene transcription by directly interacting with the transcriptional machinery, some TFs known as pioneer factors may regulate gene expression by remodeling the chromatin landscape to control transcriptional activity^9^. The accessibility of TFs to DNA is strictly controlled by the dynamic interplay between DNA methylation at CpG sites and histone modifications in a cell-type specific manner^10,11^.

Several studies have reported DNA methylation at distal enhancer regions to be implicated in gene regulation mainly by interfering with TF binding to enhancer regions^12-14^. Enhancer methylation is known to be highly dynamic and more tissue specific than promoter methylation thereby suggesting that enhancers may play a significant role contributing to cell phenotype^15-17^. As for promoters, DNA methylation at enhancers tends to be associated with transcriptional inactivity, while enhancer hypomethylation is often associated with TF binding followed by transcriptional activation^8,18^. However, the role of DNA methylation at enhancer regions and TF binding sites is still not fully understood.

We previously presented the genome-wide expression-methylation Quantitative Trait Loci (emQTL) analysis and showed that estrogen receptor (ER) positive breast tumors display disease-specific hypomethylation of enhancers carrying binding sites of ERα, FOXA1, and GATA3, suggesting an epigenetic regulation of estrogen signaling in breast cancer^19^. The two strongest and visually most apparent clusters were reported: the described above ER cluster and a cluster related to varying immune infiltration. Here, we expand our analysis to include more patient samples and use formal biclustering methods to characterize novel biclusters of emQTL associations. We discover a proliferation-related bicluster in breast cancer characterized by hypomethylation at enhancers carrying transcription factor binding sites (TFBS) of proliferation-driving TFs in ER negative tumors. The identified CpGs and genes were found enriched in enhancer regions and to be connected through chromatin loops, thereby indicating that proliferation in breast cancer is under epigenetic regulation.

## Results

### Expanded expression-methylation Quantitative Trait Loci (emQTL) analysis

Genome-wide correlations between the expression levels of all genes and the level of DNA methylation at all CpGs was performed using a similar approach to the one described by Fleischer, Tekpli et al.^19^ using a larger discovery cohort (OSL2 breast cancer cohort, n=277; Supp. Figure 1, see Material and Methods). Pearson’s correlations between CpGs with an interquartile range of more than 0.1 (*n* = 182,620) were tested against all genes (*n* = 18,586) for non-zero correlations. We identified 16,193,303 significant CpG-gene associations (Bonferroni corrected *p*-values < 0.05) from which 10,264,807 (63.4%) were validated in the independent The Cancer Genome Atlas (TCGA) breast cancer cohort (BRCA, n=558). The validated associations involved the expression level of 6803 genes and methylation level of 64,439 CpGs. To focus on hub associations, the emQTL CpGs and genes with less than five associations were filtered out. The remaining CpGs (*n* = 44,263) and genes (*n* = 4904) with associations after filtering were included in downstream analyses. A significant correlation between the expression of a gene and methylation at a CpG is hereafter referred to as an emQTL.

In order to identify emQTLs with similar biological features, we grouped the emQTLs using Spectral co-clustering^20^ of the inverse correlation coefficients values (correlation coefficient*-1; see Discussion) obtained from the genome-wide emQTL analysis. For comparison, biclustering of the absolute correlation coefficient values were also performed (Supp. Table 2a; see Discussion). Spectral co-clustering allows all emQTL-CpGs and genes included in the analysis to be assigned to biclusters while simultaneously allowing clustering of columns and rows, both of which have been considerable limiting factors with the previous approach.

To determine the optimal number of biclusters for the spectral co-clustering algorithm, a mean square residual (MSR) score^21^ was estimated when the number of biclusters was set to be a number between 2 and 20. A lower MSR score is associated with a stronger coherence exhibited by the biclusters and thereby indicates better biclustering. In order to obtain as many informative and biologically distinct emQTL biclusters we plotted the average MSR scores as a function of number of biclusters, and selected the elbow of the plot to be the number of biclusters (Fig. 1a). In this way, five biclusters were defined (Fig. 1b, Supp. Table 1a-b): Bicluster 1 (8641 CpGs and 1085 genes), Bicluster 2 (9398 CpGs and 870 genes), Bicluster 3 (6910 CpGs and 936 genes), Bicluster 4 (10 564 CpGs and 1087 genes) and Bicluster 5 (8750 CpGs and 926 genes).

**Figure 1.**
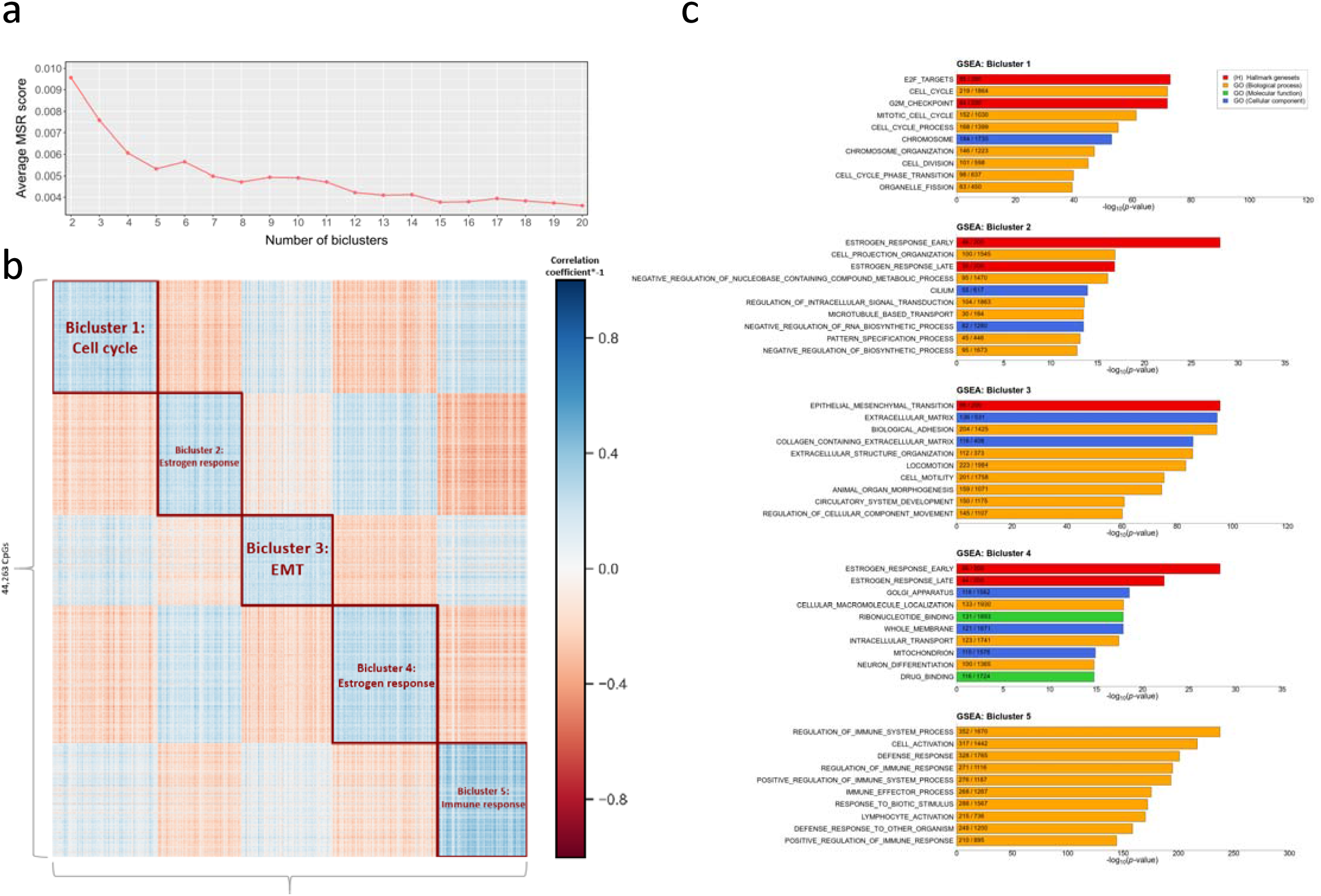
Identification and characterization of the emQTL biclusters. (**a**) Line chart showing the average MSR score for the biclusters obtained by spectral co-clustering of the inverse correlation coefficients obtained form the initial emQTL analysis OSL2 when the number of biclusters *k* were set to be a number between 2 and 20. (**b**) Heatmap displaying the five biclusters identified by spectral co-clustering when *k* was set to be 5. The biclusters will have a higher average value than the other rows and columns. Columns represents genes (*n* = 4904) and rows (*n* = 44,263) represents CpGs. Blue points indicate strong negative correlations between the variables while red points represent positive correlations. White points indicate little or no correlation. (**c**) GSEA of the genes in Bicluster 1 (*n* = 1085), Bicluster 2 (*n* = 870), Bicluster 3 (*n* = 936), Bicluster 4 (*n* = 1087) and Bicluster 5 (*n* = 926) using gene sets obtained from the MSigDB (H and C5 gene set collections). The length of the bars represents the log-transformed Benjamini-Hochberg corrected *p*-values obtained by hypergeometric distribution. Red bars indicate Hallmark gene sets while GO biological process, GO molecular function and GO cellular compartment GO gene sub-collections are colored in orange, green and blue, respectively. Overlap between the gene list of the bicluster and each MSigDB gene set is annotated within each bar.

To elucidate the biological role of the biclusters, gene set enrichment analysis (GSEA) was performed based on the genes of each bicluster using the Hallmark (H) and gene ontology (GO; C5) gene set collections (Fig. 1c and Supp. Table 1c) obtained from the Molecular Signatures Database (MSigDB^22^). As expected, we rediscovered the estrogen-(Biclusters 2/4) and immune cluster (Bicluster 5) first described by Fleischer, Tekpli et al.^19^. The majority of their immune cluster genes (94.5%) and CpGs (53.5%) were found in the newly discovered immune bicluster and the same was true for the estrogen cluster genes (53.9%) and CpGs (56.7%).

In addition to rediscovering the immune- and estrogen clusters, we now identify two novel biclusters with distinct biological functions: cell cycle regulation (Bicluster 1) and epithelial-mesenchymal transition, extracellular matrix (ECM) and cell locomotion (Bicluster 3) as shown in Fig. 1c. To confirm that the identified biclusters were not artifacts of the selected seed parameter used by the spectral co-clustering algorithm, we performed a permutation test (100 permutations) using random seeds and comparing the results with the biclustering output from the initial biclustering analysis. For each run we performed GSEA for the gene list of each bicluster identified to define which biological functions they were related. We then calculated how many times the CpGs and genes from the initial biclustering analysis described above were found in biclusters with similar biological functions. The biclusters were found to be highly stable as only 15 genes and 113 CpGs were found less than 70% of the times within a bicluster of similar biological characteristic (Supp. Figure 2a-b). We hypothesize that these CpGs and genes may represent interactions between the gene regulatory networks represented in each bicluster. Nevertheless, the biclusters were found to be stable and the inclusion of these genes and CpGs will have a negligible effect on the analyses due to their small count.

**Figure 2.**
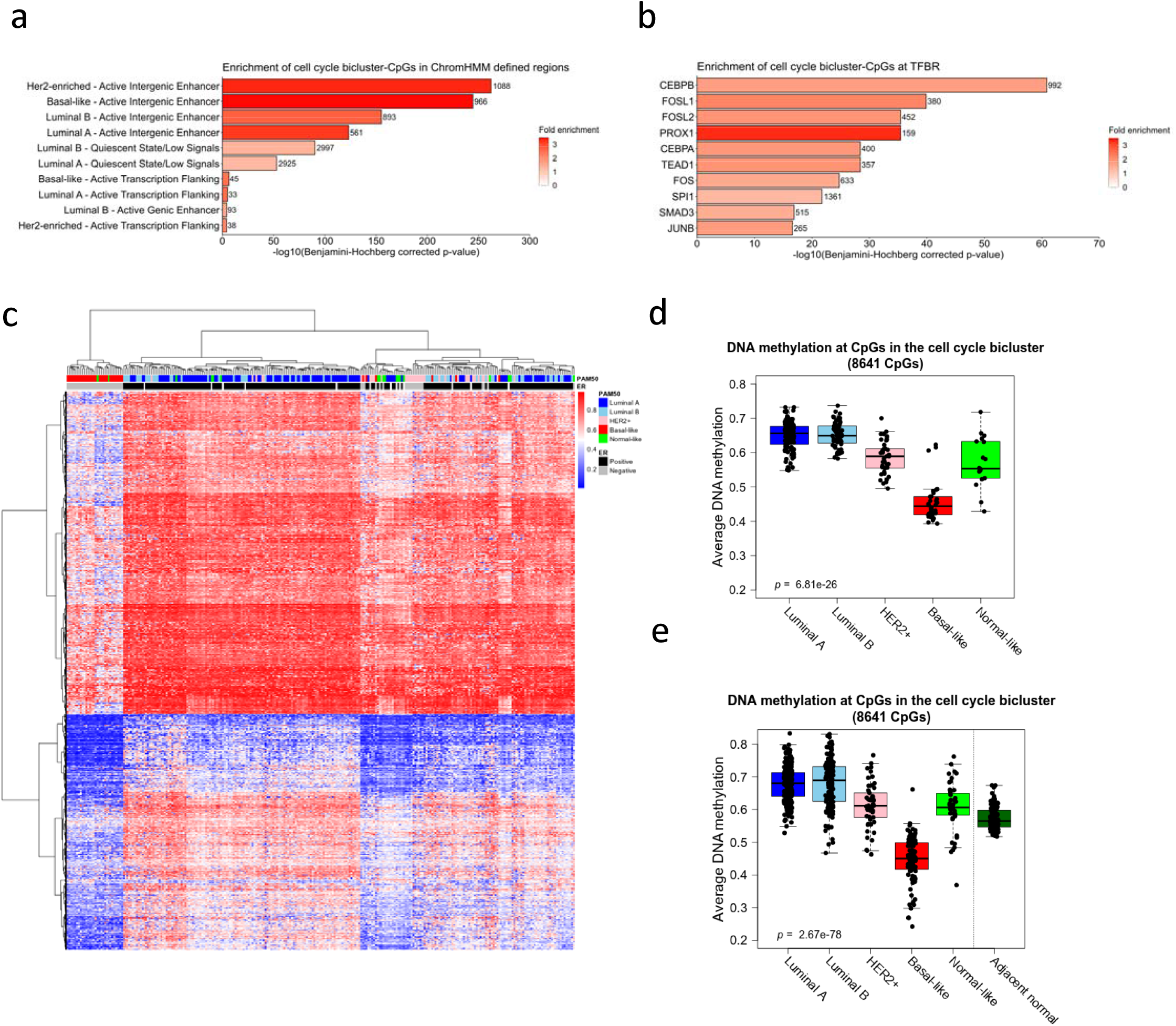
Functional characterization of the emQTL-CpGs in the cell cycle bicluster. (**a**) Bar plot showing enrichment of the cell cycle bicluster-CpGs in ChomHMM^23^ defined genomic regions by subtype. The length of the bars represents the log-transformed Benjamini-Hochberg corrected *p*-values. The color gradient of the bars represents fold enrichment in which a red color indicates FE close to 3.5 while white bars are genomic regions by subtype with FE close to 0. An enrichment was considered to be significant if the BH-corrected *p*-value was less than 0.05. (**b**) Bar plot representing enrichment of the cell cycle bicluster-CpGs at the binding site of specific TFs according to UniBind^24^. Bar length display the log-transformed BH-corrected *p*-value obtained by hypergeometric testing for each TF. Red color indicates FE close to 3.5 while a white color indicates FE close to 0. (**c**) Unsupervised hierarchical clustering of DNA methylation levels of the 8641 cell cycle bicluster-CpGs for the tumor samples in OSL2 with PAM50 status available (*n*=272). Rows represents CpGs and columns represent histopathological features including PAM50 subtype and ER status of the tumor samples. Red points indicate methylated CpGs while blue points represent unmethylated CpGs. Boxplots showing the average DNA methylation of the cell cycle bicluster-CpGs in (**d**) OSL2 (*n*=272) and (**e**) TCGA (*n*=562) by PAM50 subtype. For TGCA, DNA methylation for adjacent normal samples are also included (n=97). Kruskal-Wallis test *p*-values are denoted in the lower left corner.

### Enhancer methylation, TF binding and a proliferative phenotype of human breast tumors

To understand the functional link between DNA methylation at CpG sites and expression of genes in the cell cycle bicluster (Bicluster 1), we first aimed to characterize the functional role of the CpGs using ChromHMM segmentation data based on ChIP-seq of several histone modifications from breast cancer cell lines representing distinct subtypes^23^. According to ChromHMM, CpGs in the cell cycle bicluster were significantly enriched in active intergenic enhancer regions of breast tumors across all subtypes (Fig. 2a, Supp. Table 1d). Moreover, we found 46 % of the CpGs to overlap with at least one active intergenic enhancer region of another subtype (Supp. >Figure 3) which therefore suggests that we are discovering mechanisms regulating genes associated with proliferation that are both subtype specific as well as common across subtypes.

**Figure 3.**
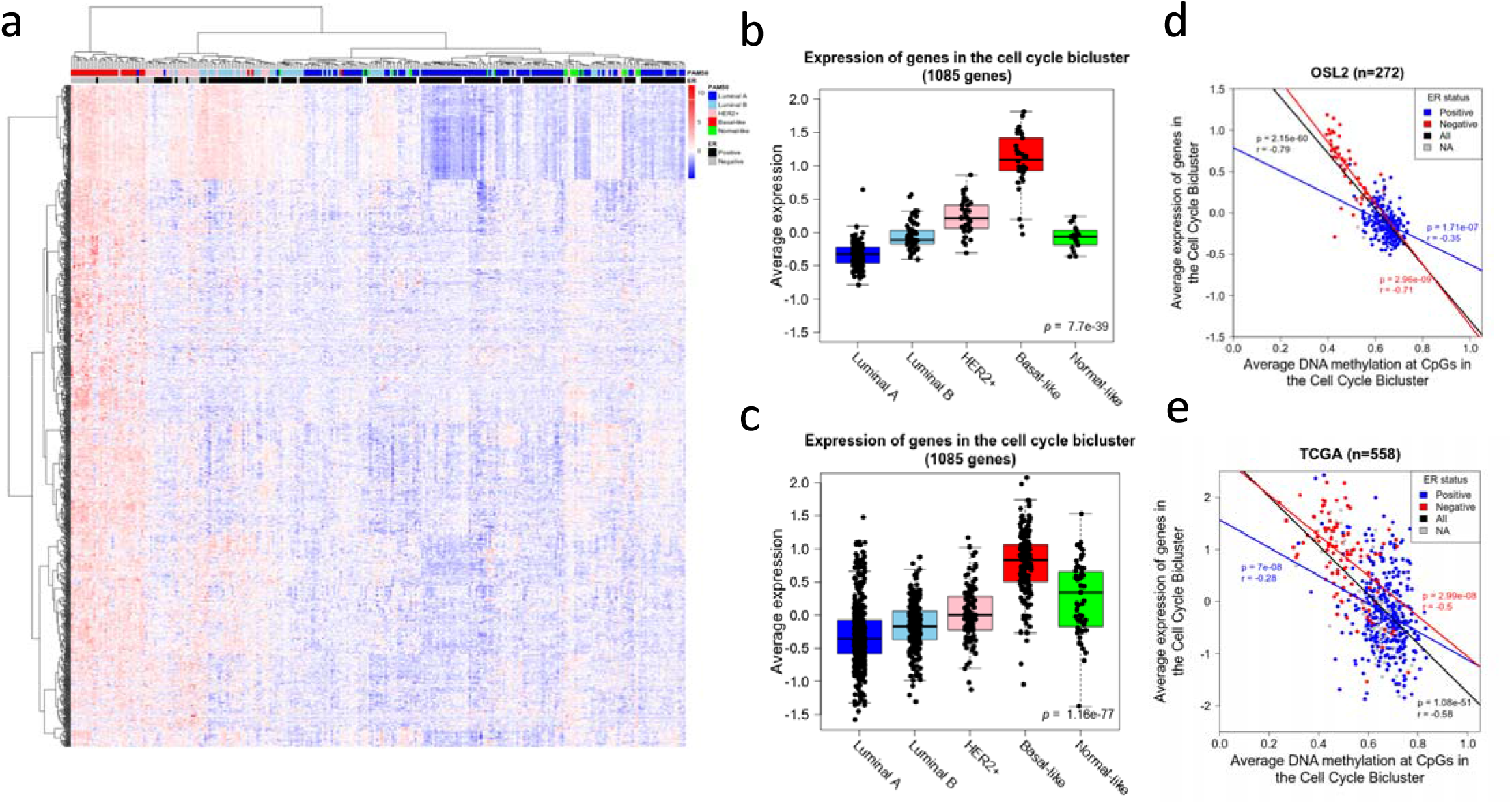
Expression of genes in the cell cycle bicluster. (**a**) Unsupervised clustering of the expression levels of the 1085 genes in the cell cycle bicluster for the tumor sin the OSL2 breast cancer cohort (*n* = 272). Rows represents genes and columns represents samples annotated with histopathological features including PAM50 subtype and ER status. Red color indicates high expression levels and blue color indicates low. Boxplots showing the average expression of genes in the cell cycle bicluster in the (**b**) OSL2 (*n* = 272) and (**c**) TCGA (*n* = 981) breast cancer cohorts. Kruskal-Wallis test *p*-values are denoted. Scatterplots showing the association between average DNA methylation of the cell cycle bicluster-CpGs versus average expression of the genes contained within the same bicluster by ER status in the OSL2-(**d**) and TCGA (**e**) breast cancer cohorts. Pearson correlation coefficients and *p*-values are denoted and colored by ER status.

Having found the cell cycle bicluster-CpGs to be enriched in intergenic enhancer regions, we next sought to identify transcription factor binding regions (TFBR) significantly overlapping with cell cycle bicluster-CpGs. We used TF-DNA interaction data obtained from UniBind^24^, which is a database storing direct TF-DNA interactions for 231 unique human TFs obtained from 1983 ChIP-seq datasets performed on 315 different cell lines and tissues. The cell cycle bicluster-CpGs were found enriched in the binding region of several TFs previously described to regulate proliferation in breast cancers including CEBP-β^25^ and several of the FOS family of proteins including FOS^26^, FOSL1^27,28^, and FOSL2^29^ (Fig. 2b, Supp. Table 1e). The TFBSs of these TFs stored in UniBind are based on ChIP-seq data from breast cancer cell lines among others. While the CEBP-β transcription factor binding sites (TFBS) have been mapped by ChIP-seq in both the estrogen receptor positive (ER+; MCF7) and ER negative (ER-; SUM159 and MDA-MB-231) breast cancer cell lines, FOSL1 and FOSL2 TFBS have been mapped in the ER-BT549 and ER+ MCF7 breast cancer cell lines by ChIP-seq respectively.

We further investigated the level of DNA methylation of cell cycle bicluster-CpGs in regard to histopathological features including ER status and PAM50 subtype. Unsupervised hierarchical clustering of the DNA methylation level of the cell cycle bicluster CpGs (n=8641) was clearly associated to the breast cancer subtypes (Fig. 2c), and the CpGs in this bicluster showed lower levels of methylation in the Basal-like, Her2-enriched and Normal-like tumors in both the OSL2 (Fig. 2d) and TCGA (Fig. 2e) breast cancer cohorts. Moreover, DNA methylation at the cell cycle bicluster-CpGs in TF binding regions of the top six most enriched TFs was found lower in the Basal-like and Her2-enriched breast tumors (Supp. Figure 4a-f). Altogether, these results show that CpGs in the cell cycle bicluster are enriched for enhancer regions overlapping TFBR of TFs associated with proliferation such as CEBP-β, FOSL1, and FOSL2. Moreover, their TFBR are found to be less methylated in Basal-like and Her2-enriched tumors.

**Figure 4.**
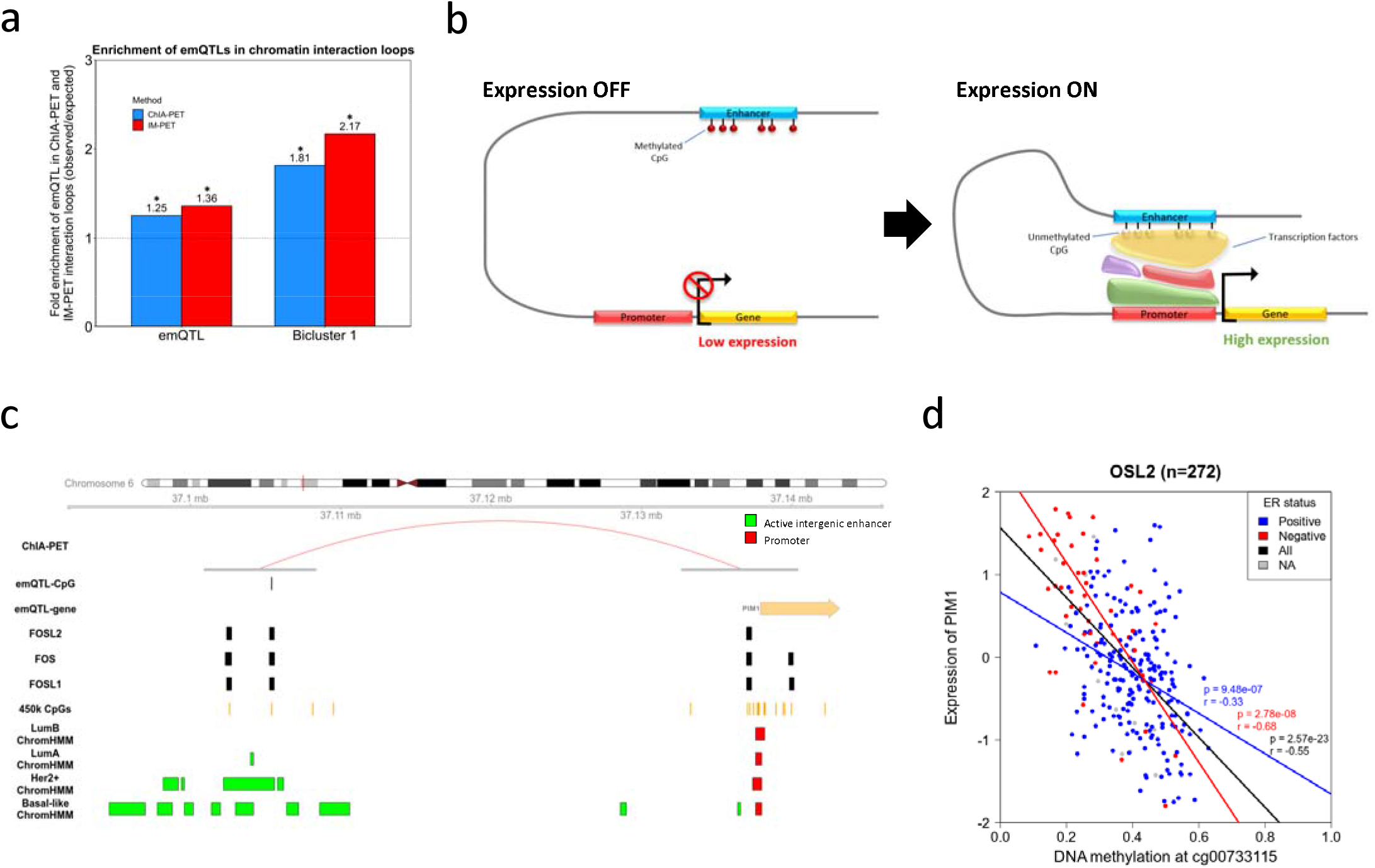
DNA methylation at enhancers facilitates target gene expression through enhancer-promoter interactions. (**a**) Bar plot showing the enrichment of emQTL-CpGs in ChIA-PET Pol2 loops and IM-PET loops for the ER+ MCF7 and ER-HCC1954 breast cancer cell lines, respectively. Bar height represents the enrichment level measured as the ratio between the frequency of emQTLs (CpG-gene pairs) found in the head and tail of a loop over the expected frequency if such overlaps were to occur at random. Enrichments that are statistically significant (hypergeometric test, Benjamini-Hochberg corrected *p*-value < 0.05) are marked with an asterisk. (**b**) Enhancers regulating proliferation-associated genes are hypomethylated which allows TF binding and transcriptional activation of the enhancer target gene through physical enhancer-promoter interaction by chromatin looping. (**c**) Example of a potential proliferation-promoting alteration in which the CpG (cg00733115) have been found in one foot of a ChIA-PET Pol2 loop (red arc) and a gene associated with proliferation (PIM1) is found in the other. Annotations for active intergenic enhancer regions and active promoters according to ChromHMM^23^ that are conserved across the cell lines of similar subtype are shown in green and blue color respectively by breast cancer subtype. The binding sites of FOS, FOSL1/2 are also shown. (**d**) Scatterplot showing the association between DNA methylation at the emQTL-CpG cg00733115 and its associated gene (PIM1) by ER status. Pearson’s correlation coefficients and *p-*values are colored by ER status.

One of the most known markers of cell proliferation is the MKI67 gene that is a non-histone nuclear protein expressed during the active phase of cell cycle^30^. To assess the link between DNA methylation at the cell cycle bicluster CpGs and proliferation we correlated the average DNA methylation levels of the cell cycle bicluster CpGs with the expression of MKI67 and found a significant negative association within the ER negative tumors and for all breast tumors (Supp. Figure 5a-b). Interestingly, we also find MKI67 to reside within the cell cycle bicluster (Supp. Table 1b). This suggests a possible link between DNA methylation at the cell cycle bicluster-CpGs and proliferation.

**Figure 5.**
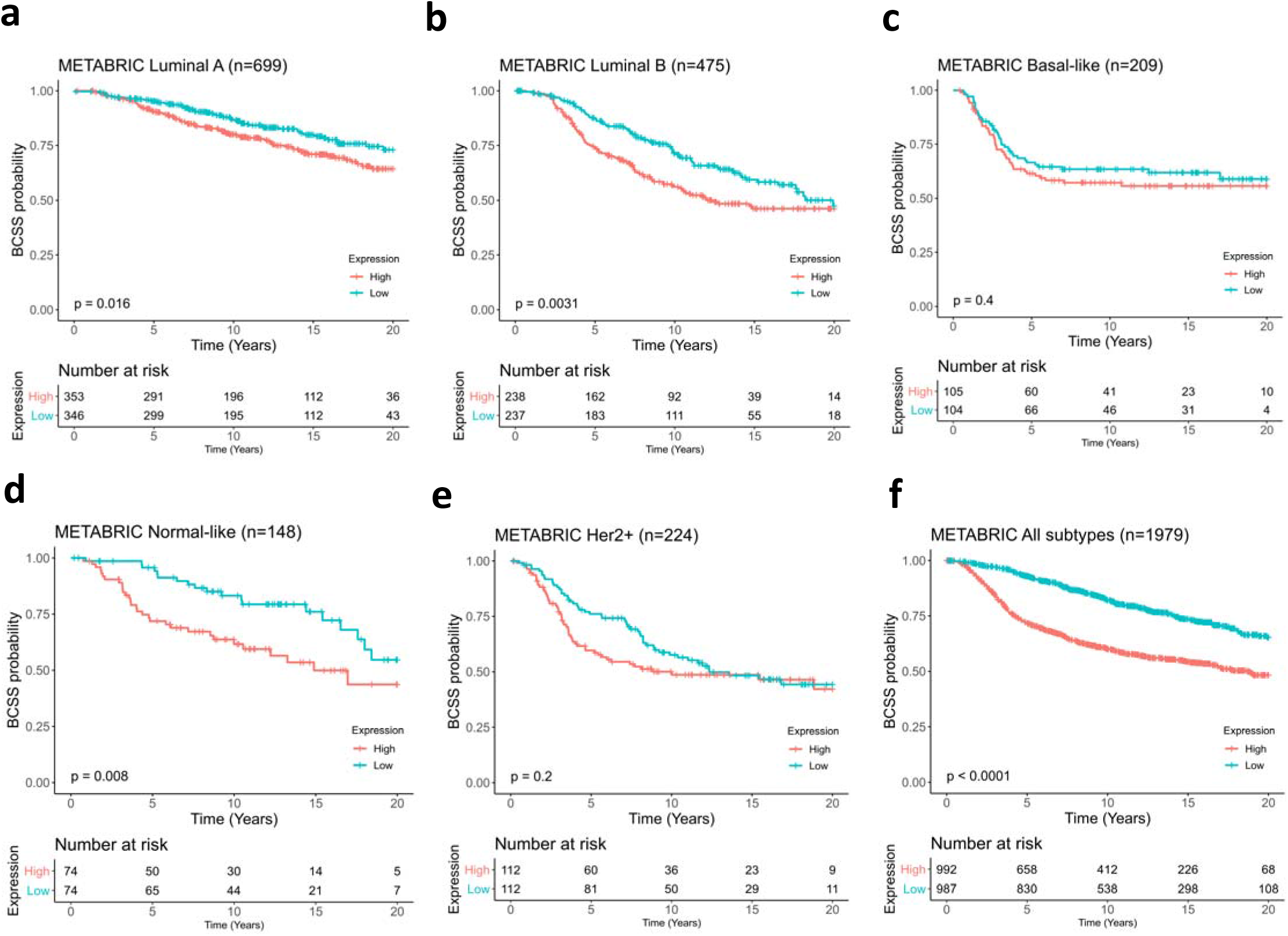
The expression levels of genes in the cell cycle bicluster is associated with prognosis. Kaplan-Meier survival curves for the cell cycle bicluster in METABRIC cohort, for Luminal A (**a**), Luminal B (**b**), Basal-like (**c**), Normal-like (**d**), Her2-enriched (**e**) and all breast cancer subtypes (**f**). Tumors were divided into two groups based on the median expression of genes in the cell cycle bicluster. *P*-values obtained by log-rank test are denoted.

To assess the link between DNA methylation (at enhancers and TF binding regions) and gene expression in the cell cycle bicluster, we performed unsupervised clustering of the expression of these genes and observed that expression was higher in the subtypes known to have higher proliferation rates (Fig. 3a). Basal-like tumors showed the highest expression, followed by Her2-enriched and Normal-like, and Luminal A showed the lowest expression (OSL2 breast cancer cohort; Fig 3b). These observations were consistent with the data from the TCGA breast cancer cohort (Fig. 3c). Next, we performed correlation analysis between the average methylation and average expression of CpGs and genes in the cell cycle bicluster. As expected, we observed a strong negative correlation between DNA methylation and gene expression (p-value 2.15e-60 and 1.08e-51; r-value = −0.79, −0.58 in OSL2 and TCGA, respectively), largely driven by the differences between ER positive and ER negative tumors (Fig 3d-e, black regression line). We also observed a significant (and validated) correlation between methylation and expression when performing the analysis separately within ER positive and ER negative (Fig 3d-e, blue and red regression line, respectively); however, the correlation was stronger within the ER negative tumors. Taken together, these results show a statistically significant association between enhancer methylation and expression of proliferation-related genes, and that ER negative breast tumors have low methylation at enhancers potentially driving proliferation. Varying degree of enhancer methylation may be related to the proliferative potential in ER negative tumors.

To investigate the functional relationship between DNA methylation and gene expression in the cell cycle bicluster we assessed the extent to which CpGs within this bicluster were located nearby (±10 kb) any of the genes contained within the same bicluster. We found that 36% of the genes in the cell cycle bicluster were located nearby at least one CpG in the same bicluster suggesting that many genes in the cell cycle bicluster may be locally regulated by DNA methylation in enhancer regions at TFBR of the enriched TFs including CEBP-β, FOSL1, and FOSL2.

Enhancers can promote gene expression of distant genes by interacting with promoter regions of their associated genes through chromatin loop formation^7,8^. Chromatin Interaction Analysis by Paired-End Tag sequencing (ChIA-PET) and Integrated Methods for Predicting Enhancer Targets (IM-PET) are methods used to identify such physical interactions on a genome-wide scale^31,32^. To more specifically assess the link between DNA methylation at enhancers and expression of their target genes, we obtained ChIA-PET Pol2 data^31^ from MCF7-(ER positive) and IM-PET interaction^32^ from the HCC1954 (ER negative) breast cancer cell lines, thereby allowing us to identify interactions between genomic loci that are located in proximity in 3D but may be separated by long distances. We found that the CpGs in the cell cycle bicluster in emQTL with cell cycle bicluster genes were significantly enriched in chromatin interaction loops as defined by ChIA-PET and in IM-PET datasets (hypergeometric test *p*-value = 5.097 x 10^−3^ and 4.74 x 10^−4^ respectively, Fig. 4a). Altogether, 59 CpGs were experimentally confirmed by ChIA-PET to form physical interactions with 39 unique genes in the cell cycle bicluster (Supp. Table 1f), and 22 unique emQTL CpG-gene loops were confirmed by IM-PET (Supp. Table 1g). Altogether, these results suggest that emQTLs represent direct regulatory links between DNA methylation at enhancer regions targeted by proliferation associated TFs and the expression of the cell cycle bicluster-genes (Figure 4b).

### Identification of potential key drivers of proliferative signaling in breast cancer

Knowing that enhancer methylation at regions of TF binding is associated with the regulation of expression of genes linked to proliferation in the cell cycle bicluster and knowing which TFs that are involved, we may refine the emQTL approach to more efficiently identify potential key drivers of carcinogenic signaling. The refined approach has several steps for identification of CpG-gene pairs: (1) The CpG-gene pair must be on opposite sides of chromatin loops defined by ChIA-PET^31^ and/or IM-PET^32^ loops. (2) A CpG must be in enhancers according to ChromHMM segmentation^23^ of either subtype. (3) The CpG must be in the binding region of the top enriched TFs as defined by UniBind^24^. Lastly, (4) the gene must be a part of a curated gene set associated with proliferation. Altogether, we identified 53 strong candidates as potential proliferation-promoting alterations in DNA methylation in which the majority display strong negative correlations between DNA methylation and gene expression (Table 1, Supp. Table 1h). Figure 4c shows an example of a potential proliferation-promoting emQTL in which the CpG (cg00733115) is found experimentally to interact with the Pim-1 Proto Oncogene, Serine/Threonine Kinase (*PIM1*) gene found in the GO_CELL_CYCLE gene set from the MSigDB^22^. The CpG itself is located in a region with high abundance of active intergenic enhancer chromatin marks in Basal-like, Her2-enriched, and Luminal A subtype according to ChromHMM^23^. Moreover, the CpG overlaps with the binding region of several members of the FOS family proteins including the FOS, FOSL1/2 TFs. The TF binding region of FOS as indicated in Figure 4c are obtained from the breast epithelial MCF10A cell line. A negative correlation was observed between DNA methylation of cg00733115 and the expression of *PIM1* in both ER negative and ER positive tumors (Fig. 4d). These results suggest that we identify strong potential proliferation-promoting alterations using the refined emQTL approach.

**Table 1.** Top potential cancer-promoting alterations identified using the refined emQTL approach. The strength of the correlations and the Bonferroni corrected *p*-value is shown for the OSL2 (n=272) and TCGA (n=558) breast cancer cohort. The table is ordered by the strength of the negative correlation in OSL2.

| emQTL ID | Pearson's correlation coefficient |  | Adjusted *p*-value |  | Method |
| --- | --- | --- | --- | --- | --- |
|  | OSL2 | TCGA | OSL2 | TCGA |  |
| cg00733115_PIM1 | -0.554 | -0.318 | 1.36E-21 | 7.38E-13 | ChIA-PET |
| cg02976539_SLC9A3R1 | -0.542 | -0.520 | 2.02E-20 | 2.69E-38 | IM-PET |
| cg18037834_KRT18 | -0.540 | -0.514 | 3.14E-20 | 2.66E-37 | ChIA-PET |
| cg15880704_PDCD4 | -0.539 | -0.288 | 3.78E-20 | 2.00E-10 | ChIA-PET |
| cg04482712_SLC9A3R1 | -0.535 | -0.541 | 7.70E-20 | 4.59E-42 | IM-PET |
| cg00484122_RHOB | -0.535 | -0.421 | 8.75E-20 | 1.15E-23 | ChIA-PET |
| cg16729850_KRT18 | -0.526 | -0.510 | 5.44E-19 | 1.41E-36 | IM-PET |
| cg21359793_KRT18 | -0.523 | -0.503 | 9.20E-19 | 2.27E-35 | IM-PET |
| cg20812370_PBX1 | -0.513 | -0.389 | 6.70E-18 | 6.52E-20 | ChIA-PET |
| cg12610744_KRT18 | -0.511 | -0.508 | 8.58E-18 | 2.98E-36 | IM-PET |

### The cell cycle bicluster associates with prognosis

To investigate the prognostic impact of the identified genes, we performed survival analysis in the METABRIC breast cancer cohort (n=1904) due to the long follow-up time. Survival analysis was performed using Kaplan-Meier estimator and log-rank tests. When stratifying tumors by PAM50 subtype and dividing the patients into two groups based on the median expression values of the genes in the cell cycle bicluster, we observe that high expression of the cell cycle bicluster-genes is associated with significantly worse prognosis within Luminal A, Luminal B and Normal-like breast tumors (Figure 5a-e, log rank *p*-value = 0.016, 0.0031, and 0.008 respectively). When performing the survival analysis independent of subtype, we observe a strong association between survival and the expression of genes in the cell cycle bicluster (Figure 5f, log rank *p*-value<0.0001).

### Rediscovery of the immune- and estrogen response related biclusters

Both the immune and the estrogen biclusters were found to overlap with the immune and estrogen-related clusters first described by Fleischer, Tekpli et al.^19^. The immune bicluster-CpGs were found enriched in close proximity to TF binding regions of several TFs involved in immune cell homeostasis such as RUNX1, FLI1 and ERG (Supp. Table 1e). DNA methylation and gene expression levels of the rediscovered immune bicluster was associated with varying degree of immune infiltration (Supp. Figure 6a-b).

**Figure 6.**
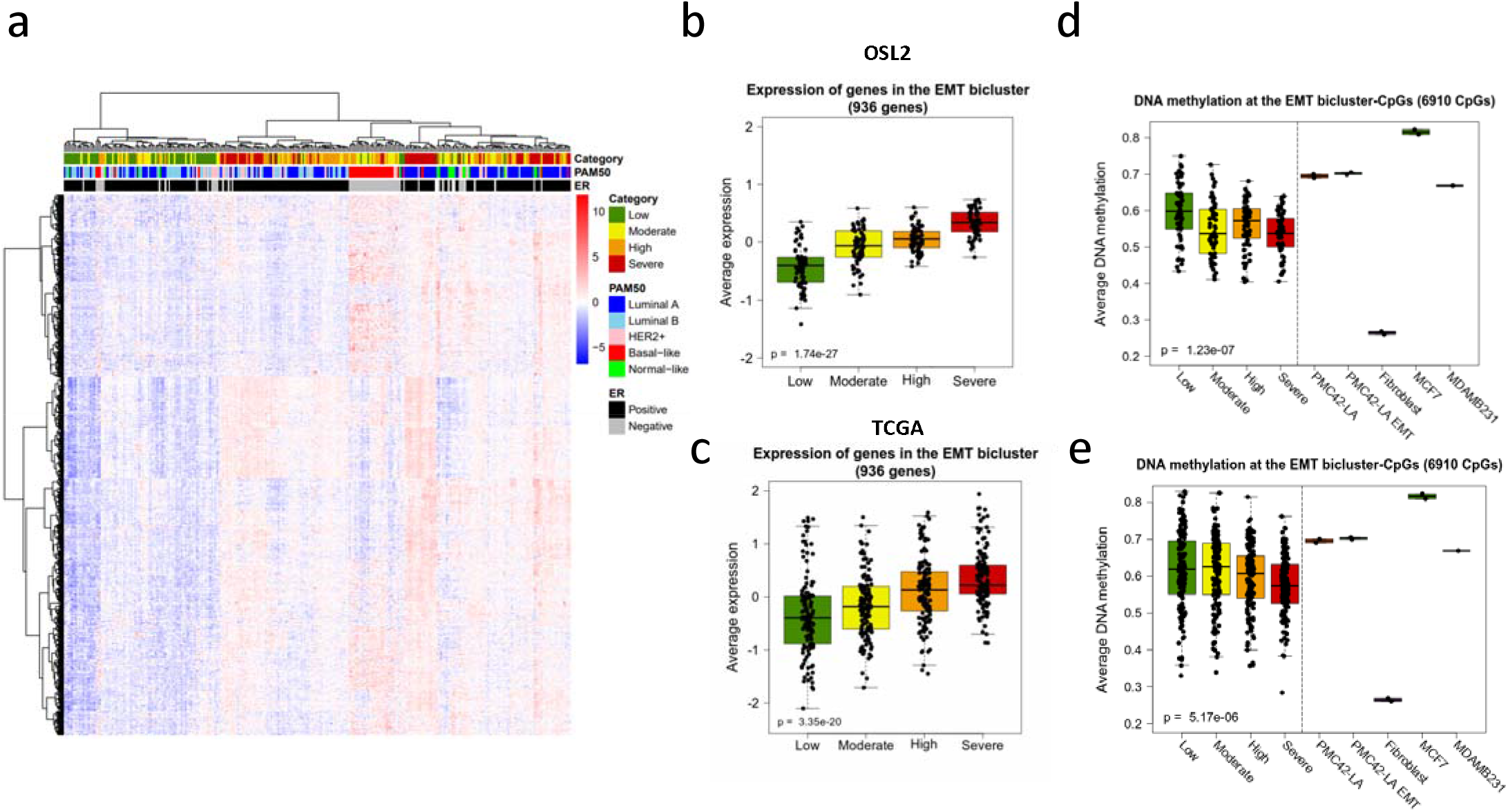
The EMT bicluster highlights an association between DNA methylation and fibroblast infiltration. (**a**) Heatmap showing the unsupervised clustering of the expression levels of the 936 genes contained within the EMT bicluster for 272 tumor samples from the OSL2 cohort. Rows represents genes and columns represent tumor samples annotated by histopathological features including PAM50 subtype and ER status. The tumor samples were divided into quartile groups based on fibroblast infiltration severity according to the relative amount of fibroblast in the tumor samples estimated by xCell^32^. Difference in expression of the EMT bicluster genes between the quartile groups are shown for the OSL2 (**b**) and TCGA (**c**) cohorts. Each quartile group consisted of 68 tumor samples in OSL2 and 139 samples in TCGA. Boxplots showing the average DNA methylation at the 6910 CpGs contained within the EMT bicluster according to fibroblast infiltration score in (**d**) OSL2 (n=272) and (**e**) TCGA (n=556). Average DNA methylation values for these CpGs for in the PMC42-LA before and after EGF induced EMT. Fibroblasts, and the ER+ MCF7 and ER-MDAMB436 breast cancer cell lines are also included. Kruskal-Wallis test *p*-values are denoted in the bottom left corner.

Contradictory to previous findings, our emQTL-CpGs and genes associated with estrogen response separated into two biclusters (Figure 1c, Supp. Table 1a-b). This is likely due to the high predominance of estrogen response-related emQTL-CpGs and genes since the spectral co-clustering algorithm favors more equally sized biclusters. The CpGs in estrogen bicluster 2 and 5 were significantly less methylated in ER+ compared to ER-tumors (Wilcoxon rank sum test, p=1.06e-18 and 1.48e-18 respectively). The estrogen-related genes in both biclusters were overexpressed in ER+ tumors compared to the ER-(Wilcoxon rank sum test, p=3.16e-24 and 9.25e-20). Moreover, the CpGs within each of the estrogen-related biclusters were enriched in enhancer regions and in genomic regions in close proximity to TF binding regions of several TFs associated with estrogen-response such as ERα, FOXA1, and GATA3 (Supp. Table 1d-e). This was observed in both estrogen biclusters which suggests that these two biclusters represent the same biological pathway. Altogether, these results are in concordance with the corresponding clusters first described by Fleischer, Tekpli et al.^19^.

### Bicluster 3 reflects varying degree of fibroblast infiltration

GSEA indicated that genes in bicluster 3 were related to processes including EMT, ECM and cell locomotion (Supp. Table 1c). Contrary to the cell cycle bicluster, genes and CpGs in the EMT bicluster (bicluster 3) to seemed to a lesser extent segregated the breast cancer patients according to the PAM50 subtypes (Fig. 6a, Supp. Figure 7a). Fibroblasts are prominent cell types of the tumor microenvironment and carry out functions related to extracellular matrix remodeling while also being able to migrate^33^. We therefore hypothesized that this bicluster was linked to fibroblast infiltration. To examine this, we estimated the relative amount of fibroblasts for each tumor sample using the xCell^34^ deconvolution tool which is based on mRNA expression. By dividing the tumors into quartile groups based on the severity of fibroblast infiltration, we found the EMT bicluster gene expression levels to be linked with fibroblast infiltration in OSL2 and TCGA (Fig. 6b-c, Kruskal-Wallis test p-value = 1.74 x 10^−27^ and 3.35 x 10^−20^, respectively) i.e., high expression of the EMT bicluster genes is linked with high fibroblast infiltration. Altogether this suggest that the expression levels of these genes may be caused by a high expression of the EMT bicluster genes in tumor-infiltrating fibroblasts rather than the cancer cells themselves.

Furthermore, we characterized the CpGs in the EMT bicluster and found them to be enriched in active intergenic enhancer regions, but to a lower extent than cell cycle bicluster CpGs (Supp. Table 1d). No significant enrichment of EMT bicluster-CpGs in emQTL with EMT bicluster genes were observed from the ChIA-PET Pol2 and IM-PET data (Supp. Figure 7b). TF enrichment analysis revealed significant enrichment of the CpGs close to TFBR of several TFs previously linked to EMT such as FOSL1^35^, TEAD1^36^, NFIC^37^ and TWIST1^38^ (Supp. Table 1e). Average DNA methylation of CpGs in the EMT bicluster was associated with varying degree of fibroblast infiltration in OSL2 and TCGA (Figure 6d-e) i.e., increasing fibroblast infiltration was associated with decreased DNA methylation. DNA methylation data for the EMT bicluster-CpGs obtained from the EMT-associated PMC42-LA breast cancer cell line featured similar methylation levels to the tumors with low fibroblast infiltration (Figure 6d-e). Moreover, DNA methylation level at the CpGs in the PMC42-LA cell line displayed no pronounced difference after EGF-induced EMT (Figure 6d-e). Noteworthy, the EMT bicluster-CpGs display low methylation levels in human mammary fibroblasts which is the inverse observation of the one for the PMC42-LA breast cancer cell line. Taken together, these results show that DNA methylation and the expression level of genes in the EMT bicluster is caused by varying degree of fibroblast infiltration.

### Cell-type specific expression of genes in the emQTL-bicluster s by scRNA-seq

Since the tumor microenvironment consist of a highly dynamic and heterogenous collection of cells, we used single-cell RNA-seq (scRNA-seq) data from 14 breast cancer patients^39^ to investigate cell type-specific expression of a subset of genes from each bicluster. For the analysis we selected out 10 genes from each bicluster showing the strongest negative correlation with an associated emQTL-CpG within the same bicluster. We found most of the genes from the cell cycle bicluster to be cancer-specific compared to other cells types such as immune cells, fibroblasts and endothelial cells which are prominent cell types of the tumor microenvironment (Figure 7a-b). Moreover, these genes were highly expressed by cancer cells from tumors classified as Her2-enriched and TNBC subtypes compared to Luminal A and Luminal B (Figure 7c-f). Altogether, this supports the hypothesis that the cell cycle bicluster genes are important regulators of proliferation in breast cancers. Similarly, to the cell cycle bicluster, the estrogen-related genes were almost exclusively expressed by cancer cells from ER+ tumors (Figure 7b-d). Contrary, the genes associated the EMT-and immune biclusters were mainly expressed by fibroblasts and immune cells respectively (Figure 7b).

**Figure 7.**
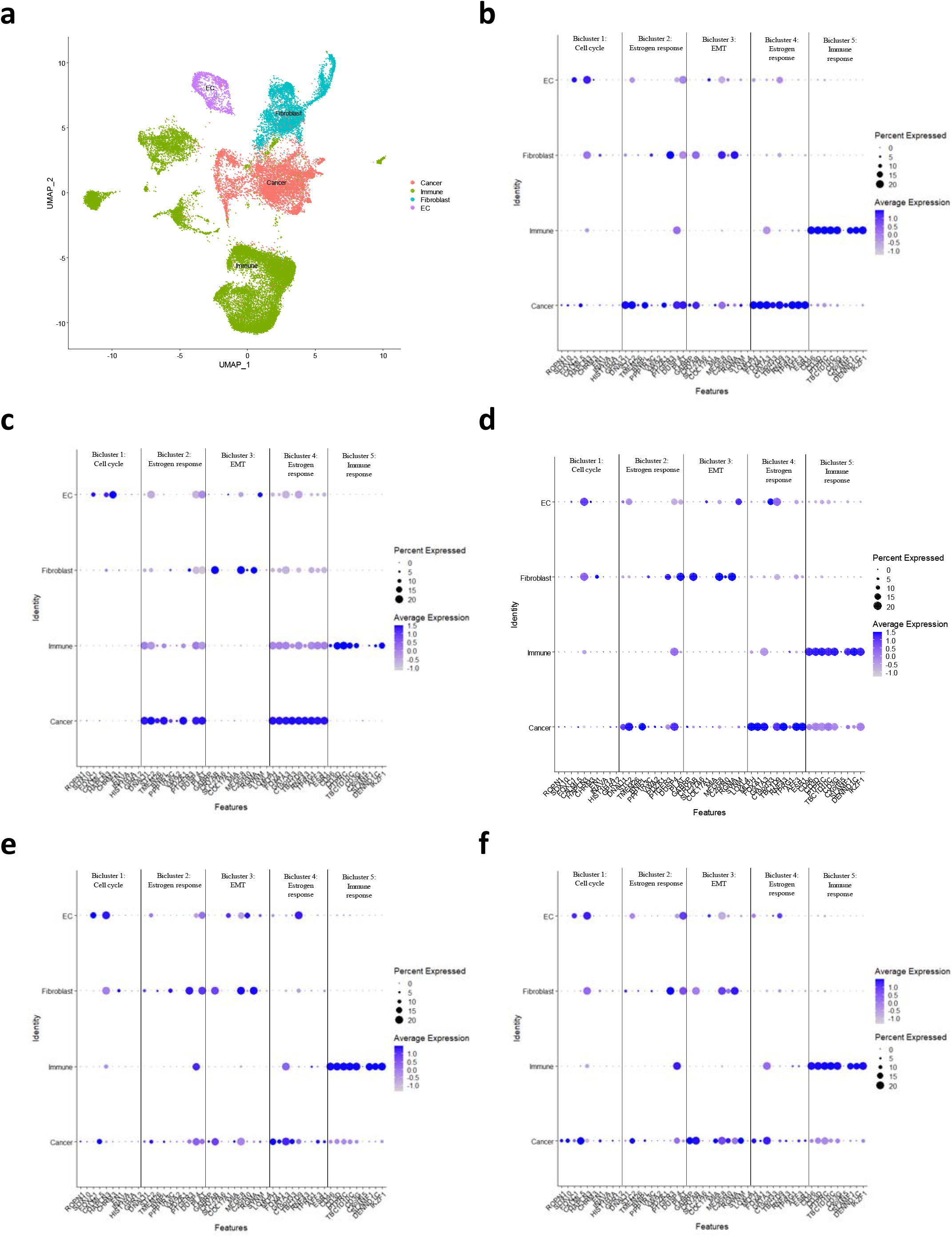
Cell-type specific expression of the emQTL-bicluster genes. Combined UMAP plot for all 14 breast cancer samples annotated by cell type (**a**). Dot plots showing the expression of the selected genes for each bicluster for **b** all patients and for patients with a (**c**) Luminal A, (**d**) Luminal B, (**e**) Her2-enriched and (**f**) TNBC. The size of the dot depicts the percentage of cells within each class and the intensity of the color shows the average expression level of each class.

## Discussion

Cancer initiation and progression involves altered proliferation rates that play an important role in breast cancer pathogenesis^40,41^. Today, little is known about how DNA methylation contributes to the proliferative phenotype of breast tumors. By performing genome-wide emQTL analysis prior to spectral co-clustering of the correlation coefficients, we identify a previously unreported gene regulatory network involved in breast cancer carcinogenesis. In ER negative breast tumors, we observe hypomethylation at enhancers carrying TFBS of key proliferation-driving TFs with concomitant high expression of proliferation-related genes in tumor cells as confirmed by scRNA-seq. We show that the identified CpGs and genes were connected through chromatin loops. Taken together, we show that proliferation in breast cancer is linked to loss of enhancer methylation and TF binding through chromatin loops. The causal effects the candidates have on the observed associations regarding the cancer phenotype will be of great interest for future studies.

Several approaches were used to identify the most optimal conditions for the biclustering algorithm in order to obtain high quality biclusters. Biclustering of the absolute correlation coefficients lead to the discovery of biclusters corresponding to the ones discussed in this paper in terms of biological characteristics such as estrogen- and immune response, cell cycle, and EMT (Supp. Figure 8, Supp. Table 2a). However, biclustering of these data were not considered optimal as it was observed a pronounced degree of spillover between all biclusters; subclusters of CpGs and genes were discovered for all biclusters with mixed functional characteristics. For instance, one subcluster of CpGs and genes in the cell cycle bicluster was strongly associated to cell cycle related functions, while the other CpG and gene subclusters were mixed with genes and CpGs related to both fibroblast infiltration and estrogen response (Supp. Figure 9-10, Supp. Table 2a). Similar observations could be made for all other biclusters. In a previous study^19^ we saw that DNA methylation was negatively associated with the activity of estrogen-related pathways and increased immune infiltration. We therefore decided to use the inverse correlation coefficients as input for the spectral co-clustering algorithm. Furthermore, the identification of potential key drivers of proliferative signaling in breast cancer at the end of the paper was not influenced by this choice as the identification pipeline was not restricted to any biclusters. The elbow method was used to determine the number of biclusters that was set to be 5, however improvement in the average MSR score was also observed for 8 and 12 biclusters. Biclusters with similar biological functions are also found when performing biclustering based on these parameters (Supp. Table 2b-c). However, increasing the number of biclusters to be a value >5 leads to excessive splitting of these biclusters representing the same biological pathways. Additional biclusters could also be discovered, however, enrichment of their genes in any gene sets were less significant. The biclusters was more mixed regarding to biological functions and showed some degree of spillover from the other biclusters.

The proliferation related CpGs were significantly enriched in active intergenic enhancer regions of all breast cancer subtypes, but most pronouncedly in Basal-like, Her2-enriched, and Luminal B tumors according to ChromHMM^23^, which are also the most proliferative subtypes. The CpGs in this bicluster were found enriched in chromatin loops in ER positive and ER negative breast cancer cell lines, thereby strengthening the hypothesis that the transcriptional network associated with proliferation could be regulated by DNA methylation independent of ER status. TF enrichment analysis showed an enrichment of the proliferation-related CpGs nearby TFBS of several TFs known to be implicated in breast cancer tumorigenesis including CEBP-β, FOSL1 and FOSL2. The CEBP family of TFs are known to be involved in regulating proliferation, and the CEBP-β member is commonly overexpressed in ER negative tumors compared to ER positive tumors and is positively associated with tumor grade^25^. Several of the Fos family TFs have also been implicated in proliferation. FOSL1 binding have previously been found enriched at enhancers of triple negative breast cancers and positively associated with proliferation in ER negative and ER positive cell lines^27^. Furthermore, FOSL2 overexpression has been linked to proliferation in the triple negative MDA-MB-231 and Her2-enriched SK-BR-2 breast cancer cell lines^29^. FOS have previously been shown to be an important regulator of proliferation in the MCF7 breast cancer cell line^26^. Here, we show that the CpGs in close proximity to the TFBS of these TFs were less methylated in the most proliferative tumor subtypes such as the Basal-like and Her2-enriched tumors. Altogether, we speculate that demethylation of the cell cycle bicluster-CpGs leads to more frequent binding of proliferation-related TFs and looping to their associated gene, thereby causing enhanced expression. The predictive and prognostic relevance of DNA methylation levels around the genomic regions binding CEBP-β, FOSL1 and FOSL2 constitute interesting regions for further investigation. At present, there is a lack of ChIP data mapping genome-wide TF-DNA interactions. Therefore, there may be other TFs as well involved in TF binding at the specified enhancers that are not included here and might also be drivers of proliferation in breast cancer.

By characterizing several aspects of the regulatory pathways associated with proliferation in breast cancers, we were able to identify potential downstream drivers of carcinogenic signaling relating to proliferation. The identified candidate gene with the strongest and most significant negative correlation was the *PIM1* gene which belongs to the Serine/Threonine protein kinase family of proteins. PIM1 is known to be implicated in the cell cycle, and knockdown experiments in triple negative breast cancer cell lines have been shown to decreased proliferation and survival^42^. Another candidate was *CDKL3* which is a *CDK3* homolog belonging to the cyclin-dependent protein kinase (CDK) family of proteins. *CDKL3* is known to be implicated in cell cycle progression from G1 to the S phase^43,44^. The methylation status of an emQTL-CpG located in a distal enhancer region was found linked to the expression of the *CDKL3* gene though chromatin looping defined by an experimentally defined ChIA-PET Pol2 loop. A previous study found CDKL3 upregulation to be associated with faster-growing HeLa cells derived from cervical cancer^45^. However, less is known about the influence of *CDKL3* upregulation on proliferation in breast cancer. Another candidate such as MUC1, which is an oncoprotein, has been linked to proliferation in breast cancer cell lines upon siRNA knockdown experiments^46^. Cyclin D1 (CCND1) is involved in progression of several cancer types including breast, lung, esophagus and bladder cancers. CCND1 is associated with proliferation by regulating the G1/S-phase transition^47^. Knockdown of *CCND1* using siRNA have been shown to decrease proliferation rates in the MCF7 breast cancer cell line^48^. Altogether, this indicates that our identified proliferation-promoting candidate genes play key roles in proliferation-related processes in breast cancer.

Previous studies have linked increased proliferation rates with prognosis in breast cancers^40,41^. Here, we report the expression of the proliferation-related genes in the cell cycle bicluster to be associated with poorer prognosis within the established breast cancer subtypes, including Luminal A and Luminal B. We are thereby identifying subgroups of patients which may benefit from more aggressive treatment, and equally importantly, we identify a subgroup of patient that may benefit from less treatment.

Fibroblasts, also known as cancer-associated fibroblasts (CAFs) in a tumor setting, are among the most abundant cell types of the tumor microenvironment involved in functions related to ECM remodeling^49,50^. They also play a key role in promoting tumorigenesis^51^. An increasing number of studies have emphasized a possible link between infiltration of CAFs and epigenetic changes in tumor cells. One of the most characterized CAF-secreted factors, TGF-β, can mediate epigenetic changes through SOX4 activation, which in turn modulates the EZH2 histone methyltransferase in cancer cells^52^. Moreover, aberrant DNA methylation can occur on a genome-wide scale in tumor cells treated with TGF-β^53,54^. Fibroblast infiltration has been associated with treatment response and metastatic potential of cancer cells^55-58^. By using the xCell^34^ deconvolution tool which is based on gene expression data, we found strong associations between fibroblast infiltration versus expression and methylation levels of genes and CpGs in the EMT bicluster in OSL2 and TCGA (Figure 6b-e). Lower DNA methylation at the EMT bicluster-CpGs was associated with higher fibroblast infiltration and fibroblasts were unmethylated at these CpGs compared to tumor tissue. The emQTL analysis highlights how DNA methylation and gene expression levels may reflect infiltration levels in the tumor microenvironment. Even though the EMT bicluster is significantly associated with fibroblast infiltration, there may be a less pronounced EMT-related signal from the tumors themselves represented in the EMT bicluster caused by fibroblast infiltration or other factors. Therefore, a more detailed study of the epigenetic effects of crosstalk between fibroblasts and tumor cells regarding the EMT bicluster would be of future interests.

In this study, we provide genome-wide evidence that DNA methylation at intergenic enhancer regions is a key regulator of proliferation in breast cancers. The CpG sites involved were proximal to TFBSs of CEBP-β, FOSL1, and FOSL2, which are TFs associated with proliferation in breast cancers. Altogether, we establish an association between DNA methylation and tumor phenotype reflecting the proliferative potential of breast cancer tumors.

## Material and methods

### Patient material

The OSL2 breast cancer cohort^59,60^ has collected material from breast cancer patients with primary operable disease (T1-T2) in several south-eastern Norwegian hospitals. Patients were included between 2006 and 2019. All patients have provided written consent for use of the material for research purposes. Clinical data including PAM50 classification and mRNA expression data can be obtained from GEO with accession number GSE58215^60^ and DNA methylation data is available at GEO with the accession number GSE84207^19^ (n=277).

The Cancer Genome Atlas Program (TCGA) breast cancer cohort has previously been described^61^. Level 3 gene expression and DNA methylation data were downloaded from the TCGA Data portal (https://tcga-data.nci.nih.gov). CpGs and genes with more than 50% missing values were excluded and the remaining missing methylation values were imputed using the pamr R package (function *pamr*.*knnimpute*) with *k* = 10. Only breast cancer tumor samples with gene expression and DNA methylation data were included for validation of emQTLs in TCGA (n=558).

The Molecular Taxonomy of Breast Cancer International Consortium (METABRIC) has previously been described^62^. METABRIC is a large gene expression cohort with long follow-up time widely used for investigation of breast cancer diseases. Gene expression data is available from the European Genome Phenome Archive (DOI: EGAS00000000083, n=1980).

### Statistical compution and bioinformatical analyses

All computational analyses were performed using the R software version 3.5.1^63^ unless otherwise specified. The emQTL analysis R code can be found at https://github.com/JorgenAnkill/emQTL.

Results were considered statistically significant if the adjusted *p*-value was < 0.05. Bar plots displaying ChromHMM and UniBind enrichment results were generated using the R package *ggplot2*^64^. Kaplan-Meier estimators and log-rank tests were performed using the *survival* R package v3.2.3 (functions *Surv* and *survfit*). Survival plots were made using the *survminer* R package (v0.4.8). Upset plot was generated using the UpSet R-package version 1.4.0^65^.

### Genome-wide cor relation analysis

The DNA methylation level of all CpGs with interquartile range of more than 0.1 (*n*=182,620) were correlated by Pearson correlation with the expression of 18,586 genes in the OSL2 breast cancer cohort resulting in more than three billion tests. CpG-gene associations with a Bonferroni corrected *p*-value of less than 0.05 (nominal *p*-value < 1.47 x 10^−11^) were considered significant. The significant CpG-gene associations in OSL2 were subsequently validated in the TCGA breast cancer cohort (*n*=558). The significant associations were considered validated if the Bonferroni corrected *p*-value was less than 0.05 (nominal *p*-value < 6.70 x 10^−11^). Of the 5,928,496 non-validated emQTL pairs, 28,523 associations could not be tested due to missing DNA methylation or expression data in TCGA. Only validated associations were included in the subsequent analyses. Probes and genes with less than 5 associations were filtered out. The remaining CpGs and genes with associations were kept in the following analyses. Prior to the analysis, gene symbols for expression data in the discovery and validation cohort were harmonized using the R package *HGNChelper* version 0.7.1 (function *checkGeneSymbols*).

### Biclustering of the emQTL correlation coefficients

The inverse correlation coefficients (r*-1) from the emQTL analysis were biclustered using Python (version 3.7.9) by applying the *SpectralCoclustering* algorithm contained within the scikit-learn library^66^. For the initial spectral co-clustering analysis, the random_state parameter was set to 0. Spectral co-clustering was performed using the inverse correlation coefficients (correlation coefficient*-1) values obtained from the OSL2 discovery cohort. Python code used for biclustering is available from https://github.com/JorgenAnkill/emQTL.

### Gene set enrichment analysis

Gene sets used for gene set enrichment analysis were downloaded from the Molecular Signatures Database v7.1 (MSigDB)^22^. Enrichment was determined by hypergeometric testing (R function *phyper*) using the H and C5 gene set collections. P-values were corrected for multiple testing using Benjamini-Hochberg procedure (R function *p*.*adjust*).

### Hierarchical clustering of DNA methylation and gene expression levels

Hierarchical clustering of the DNA methylation- and gene expression levels was performed using the R package *pheatmap* using Euclidean distance and ward.D2 cluster agglomeration method. For visualization purposes, gene expression values were centered and scaled by rows by dividing the centered rows by their standard deviations (R function *scale*).

### Genomic segmentation and annotation

ChromHMM segmentation data from cell lines representing different breast cancer subtypes were obtained from Xi et al.^23^, which included MCF7 and ZR751 (Luminal A), UACC812 and MB361 (Luminal B), HCC1954 and AU565 (Her2+), HCC1937 and MB469 (Basal-like). ChIP-seq peaks for key histone modifications including H3K4me3, H3K4me1, H3K27me3, H3K9me3, and H3K36me3 were used to predict chromatin states across the genome of the cell lines. The genomes were annotated into thirteen distinct chromatin states including: active promoter (PrAct), active promoter flanking (PrFlk), active transcription (TxAct), active transcription flanking (TxFlk), active intergenic enhancer (EhAct), active genic enhancer (EhGen), bivalent promoter (PrBiv), bivalent enhancer (EhBiv), repressive polycomb domain (RepPC), weak repressive domain (WkRep), repeat/ ZNF genes (RpZNF), heterochromatin (Htchr) and quiescent state / low signals (QsLow). Subtype specific ChromHMM annotations were made by collapsing the ChromHMM annotations from cell lines of similar subtype and keeping the ones that were common.

Enrichment of CpGs in a ChromHMM defined functional region was measured as the ratio between the frequency of cell cycle bicluster-CpGs found in a specific segment type over the frequency of CpGs from the Illumina HumanMethylation450 array found within the same segment type. P-values were obtained by hypergeometric testing with the Illumina 450k array probes as background (*n*=485,512). P-values were corrected for multiple testing using Benjamini-Hochberg procedure.

### TF enrichment analysis in UniBind defined TF binding sites

Enrichment of CpGs in TF binding sites was assessed using data obtained from the UniBind 2018^24^ database. Maps of direct TF-DNA interactions were downloaded from the UniBind website (https://unibind2018.uio.no) for prediction model PWM. The genomic positions of all CpGs from the Illumina 450k array were lifted over from hg19 to hg38 using the LiftOver webtool from UCSC genome browser (https://genome.ucsc.edu) and were extended with 150 bp upstream and downstream. Since each TF can have binding sites derived from multiple ChIP-seq experiments, we merged the TF binding sites for all ChIP-seq experiments for each TF. Enrichment of CpGs in proximity to TF binding sites was computed using hypergeometric testing (R function *phyper*) with IlluminaMethylation450 Bead Chip CpGs as background. False discovery rate was estimated by Benjamini-Hochberg correction using the R function *p*.*adjust*.

### Cell line data

Illumina 450k methylation data from the epithelial-like PMC42-LA breast cancer cell line before and after EGF-induced EMT were obtained from GEO with accession number GSE97853. Human mammary fibroblasts Illumina 450k array data was obtained at GEO with accession number GSE74877^67^. DNA methylation data for the ER positive MCF7 and ER negative MDAMBA453 cell lines were obtained from GEO with accession number GSE69118^68^ and GSE124368^69^.

### scRNA-seq data

Count matrix of single cell RNA-seq obtained from Qian et al.^39^ were analyzed using the Seurat R package version 3.2.1^70^ to obtain UMAP. In brief, the count matrix was already filtered for dying cells by the authors. It was further normalized and scaled regressing out potential confounding factors (number of UMIs, number of gene detected in cell, percentage of mitochondrial RNA). After scaling, variably expressed genes were used to construct principal components (PCs). PCs covering the highest variance in the dataset were selected based on elbow and Jackstraw plots to build the UMAP. Clusters were calculated by the *FindClusters* function with a resolution between 0.8 and 1.8, and visualized using the UMAP dimensional reduction method.

Four main cell types were identified on these UMAP, combining both the information obtained from the UMAP clustering and cell type annotation from the authors. The main cell types were immune-, cancer-, endothelial cells and fibroblasts.

### xCell analysis

The xCell^34^ algorithm was used to deconvolute the cellular composition of the tumor samples. xCell is a powerful machine learning framework trained on 64 immune and stromal cell datasets used to generate cell-type-specific enrichment scores and adjusting them to cell type proportions. The algorithm uses 10,808 genes as signatures to identify specific cell types from bulk tissue. The cell type enrichment scores were calculated for the OSL2 cohort (*n*=272) using the xCell^34^ web tool (http://xcell.ucsf.edu/). Gene names from the expression data of the OSL2 cohort were harmonized with the gene list provided by the xCell tool prior to the analysis using the HGNChelper v0.7.1 R package. Pre-calculated xCell scores for TCGA tumor samples were downloaded from http://xcell.ucsf.edu/xCell_TCGA_RSEM.txt.

### Chromatin interaction mapping

ChIA-PET data defining long-range chromatin interactions in the ER positive MCF7 breast cancer cell line was obtained from ENCODE (Accession number ENCR000CAA^31^). Only *in cis* loops were included in the analysis. An emQTL was considered to be in a ChIA-PET Pol2 loop if the CpG and transcription start site (TSS) of its associated gene were found within the genomic intervals of two opposite feet of the same loop. Enrichment of CpGs in ChIA-PET Pol2 loops were calculated using hypergeometric test (R function *phyper*) with all possible *in cis* pairs between CpGs and genes of the Illumina HumanMethylation 450 Bead ChIP array as background. Computational chromatin interactions predicted by the IM-PET algorithm for the ER negative HCC1954 breast cancer cell line was retrieved from the 4Dgenome data portal^71^. BEDTools v2.27.1^72^ was used to intersect the CpG and gene positions with the genomic intervals defining the feets of the chromatin loops for the ChIA-PET and IM-PET data. Chromatin interaction plots were made using the Gviz (v1.32.0)^73^ and GenomicRanges (v1.40.0)^74^ R packages. Genome interaction tracks were made using the R package *GenomicInteractions* (v1.22.0)^75^.

## Acknowledgements

This work was supported by grant from South-Eastern Norway Regional Health Authority (grant 2020031 to TF). We would like to acknowledge Daniel Nebdal for technical support and assistance during the project. SMB and MRA were postdoctoral fellows of the Norwegian Cancer Society (grant 711164) to VN Kristensen.

## Abbreviations

BH: Benjamini-Hochberg
CAF: Cancer-associated fibroblast
ChIA-PET: Chromatin Interaction Analysis by Paired-End Tag Sequencing
ChIP: Chromatin Immunoprecipitation Sequencing
ECM: Extracellular matrix
EMT: Epithelial-mesenchymal transition
ER: Estrogen receptor
FE: Fold enrichment
emQTL: Expression-methylation Quantitative Trait Loci Analysis
GEO: Gene Expression Omnibus
GSEA: Gene set enrichment analysis
IM-PET: Integrated Methods for Predicting Enhancer Targets
METABRIC: The Molecular Taxonomy Breast Cancer International Consortium
MSigDB: Molecular Signatures Database
PCs: Principal components
scRNA-seq: Single-cell RNA-sequencing
TCGA: The Cancer Genome Atlas
TF: Transcription factor
TFBR: Transcription factor binding region
TFBS: Transcription factor binding site
TNBC: Triple-negative breast cancer
UMAP: Uniform Manifold Approximation and Projection

## Supplementary Tables

**Supplementary Table 1**. Table of all the validated emQTL (**a**) CpGs (*n*=44,263) and (**b**) genes (*n*=4.904) obtained by spectral co-clustering of the inverse correlation coefficients from the OSL2 cohort by bicluster. Genomic locations shown in the tables are based on the hg19 genome assembly. (**c**) GSEA of the genes in Bicluster 1 (*n* = 1085), Bicluster 2 (*n* = 870), Bicluster 3 (*n* = 936), Bicluster 4 (*n* = 1087) and Bicluster 5 (*n* = 926) using the MSigDB H and C5 gene set collections. (**d**) Enrichment of CpGs in ChromHMM^23^ defined regulatory regions by bicluster. (**e**) Enrichment of CpGs in UniBind^24^ defined TFBS (±150 bp) by bicluster. Intra-bicluster in *cis* ChIA-PET Pol2 loops (**f**) and IM-PET loops (**g**) for all the five identified biclusters. (**h**) shows the identified potential proliferation-promoting candidates. In order for the proliferation-promoting candidate to be considered valid, the CpG had to be in opposite sides of ChIA-PET and IM-PET loops of a proliferation-related gene and be located in an active intergenic region of either breast cancer subtype according to ChromHMM^23^. The CpG had to be found in the TFBR of one of the significantly enriched TFs from the TF enrichment analysis for the bicluster 1-CpGs in UniBind^24^.

**Supplementary Table 2**. GSEA of the genes in each of the five biclusters identified by spectral co-clustering of the absolute correlation coefficient values obtained from the emQTL analysis in OSL2 for the 44,263 and 4904 validated emQTL-CpGs and genes respectively (**a**). GSEA of the genes in each bicluster when performing spectral co-clustering of the inverse correlation coefficients when the number of biclusters *k* were set to be 8 (**b**) or 12 (**c**).

