## Supplementary Figures for "Epigenetic alterations at distal enhancers are linked to proliferation in human breast cancer"

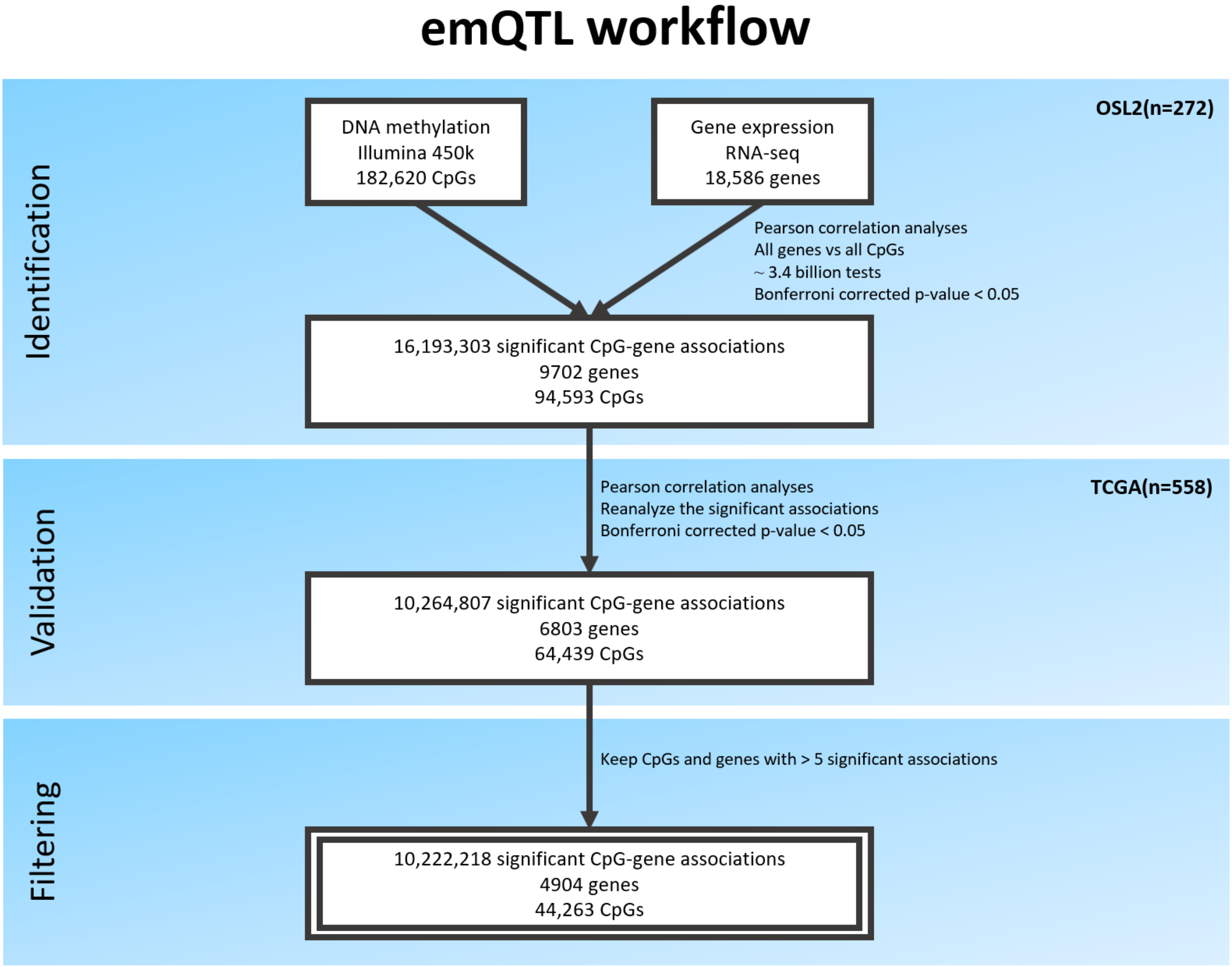


**Supplementary Figure 1. The emQTL analysis pipeline.** Flowchart showing the outline of the emQTL analysis pipeline resulting in the identification of the 10,264,807 expression-methylation Quantitative Trait Loci (emQTLs).

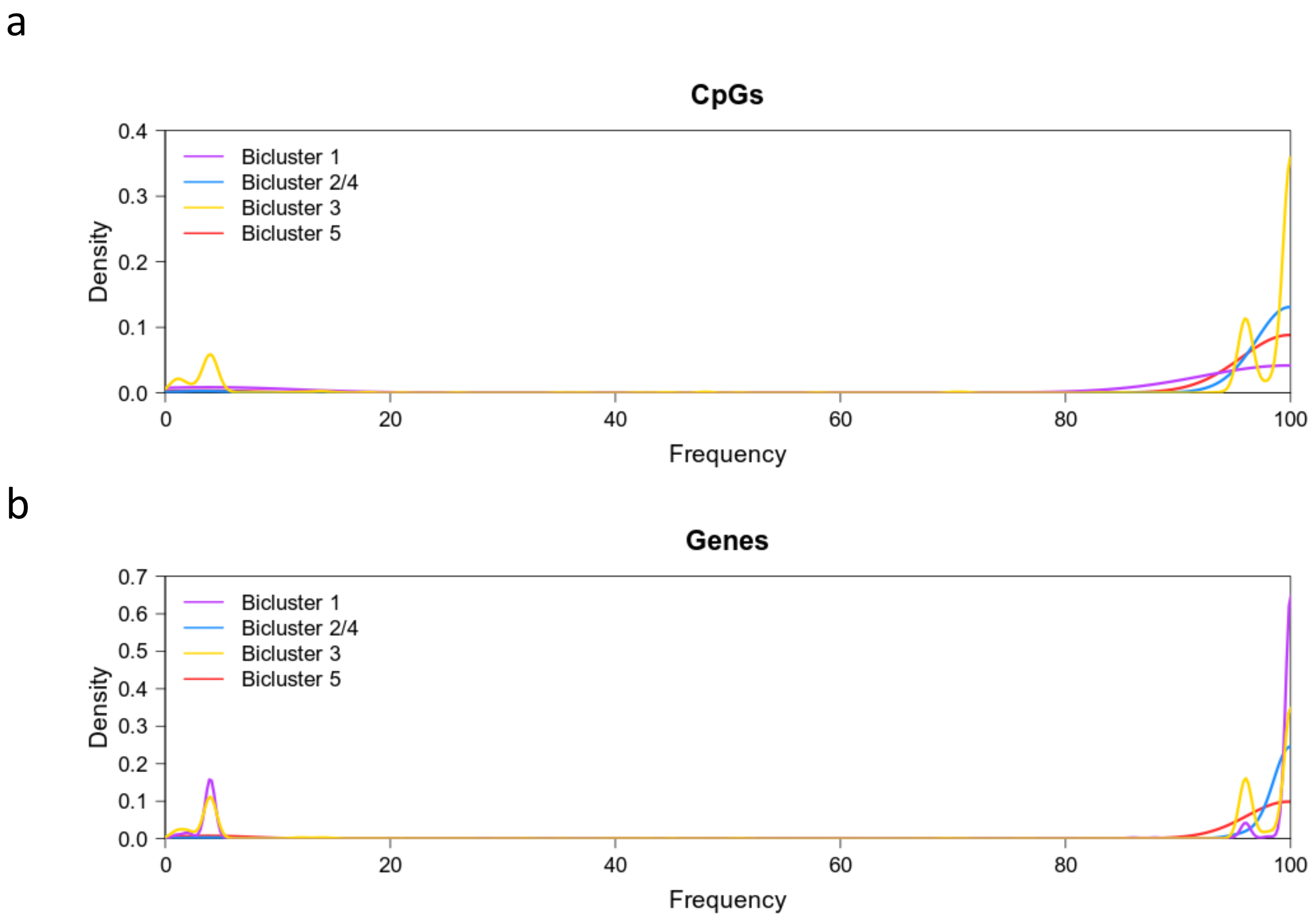


**Supplementary Figure 2. Assessment of Bicluster stability.** Density plot showing the frequency of which the emQTL probes (**a**) and emQTL genes (**b**) remains within a bicluster with similar characteristics, as the initial biclustering analysis (random_state=0), when spectral co-clustering was performed by using 100 randomly selected seeds. The two estrogen biclusters (Bicluster 2 and 4) were combined and considered as one bicluster for this analysis.

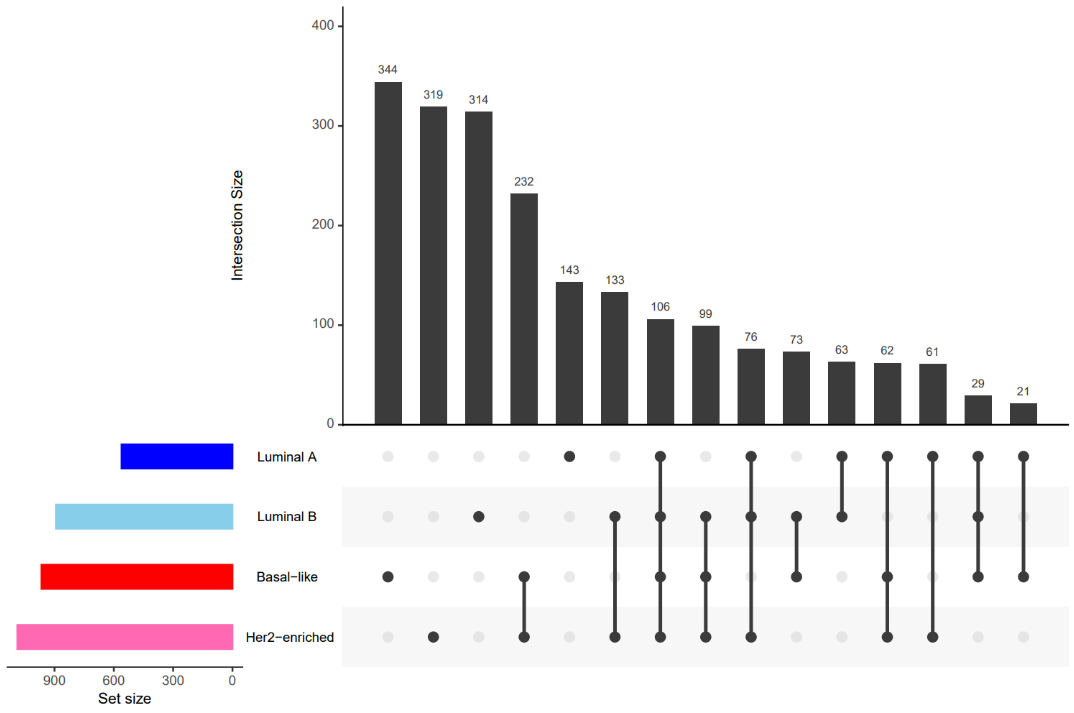


**Supplementary Figure 3.** UpSet plot showing the overlap between CpGs in the cell cycle bicluster found within ChromHMM-defined[23] active intergenic enhancer regions by breast cancer subtype.

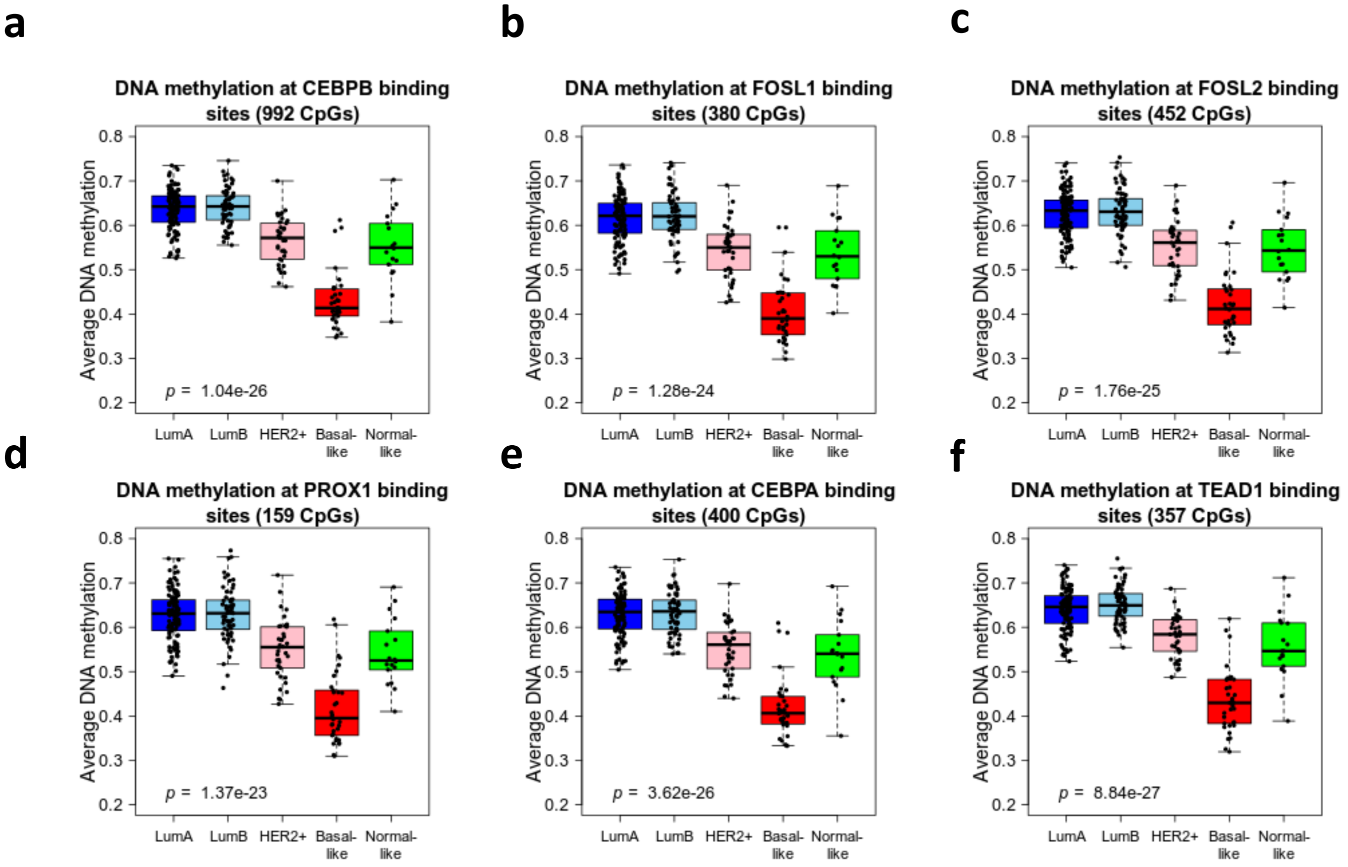


**Supplementary Figure 4. DNA methylation at TFBR in the cell cycle bicluster.** Boxplots showing the average DNA methylation of CpGs in the cell cycle bicluster at CEBP-β (**a**), FOSL1 (**b**), FOSL2 (**c**), PROX1 (**d**), CEBP-α (**e**) and TEAD1 (**f**) TFBR defined according to UniBind[24]. Boxplots represents the average DNA methylation of these CpGs in Luminal A (n=120), Luminal B (n=63), Her2-enriched (n=37), Basal-like (n=34) and Normal-like tumors (n=18) for the OSL2 breast cancer cohort. Kruskal-Wallis test *p*-values are denoted in the bottom left corner of each plot.

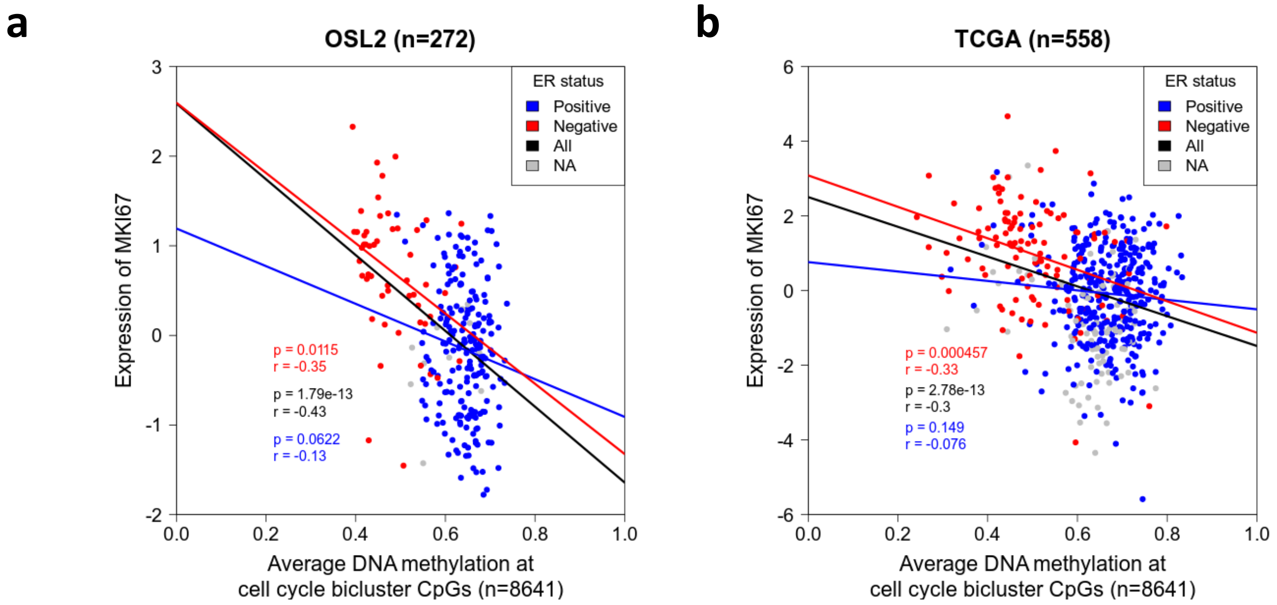


**Supplementary Figure 5.** Scatterplot showing the association between expression of the proliferation marker gene MKI67 and the average DNA methylation at the cell cycle bicluster CpGs colored by ER status in OSL2 (**a**) and TCGA (**b**).

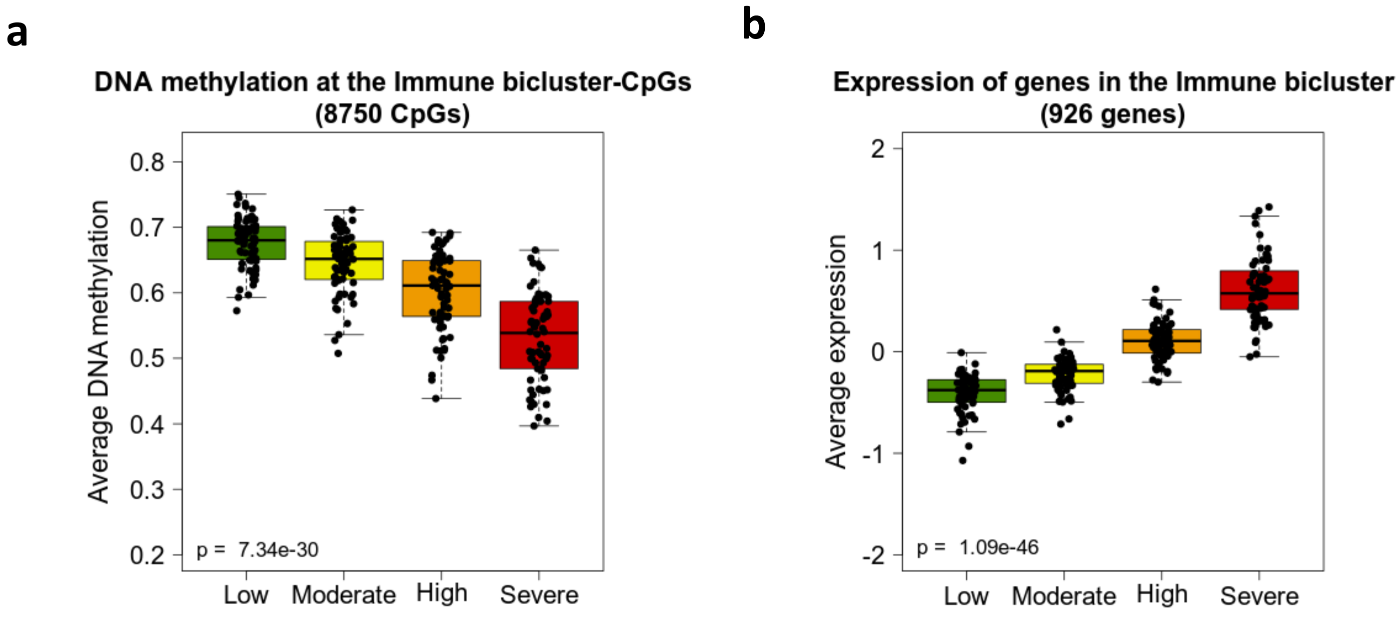


**Supplementary Figure 6. The immune bicluster is linked to varying degree of immune infiltration.** Boxplots showing the average DNA methylation of the 8750 CpGs (**a**) and expression of the 926 genes (**b**) contained within the immune bicluster according to the xCell derived immune infiltration score in OSL2 (n=272). Tumor samples were divided into quartile groups based on the severity of immune infiltration. Kruskal-Wallis test *p*-values are denoted in the bottom left corner.

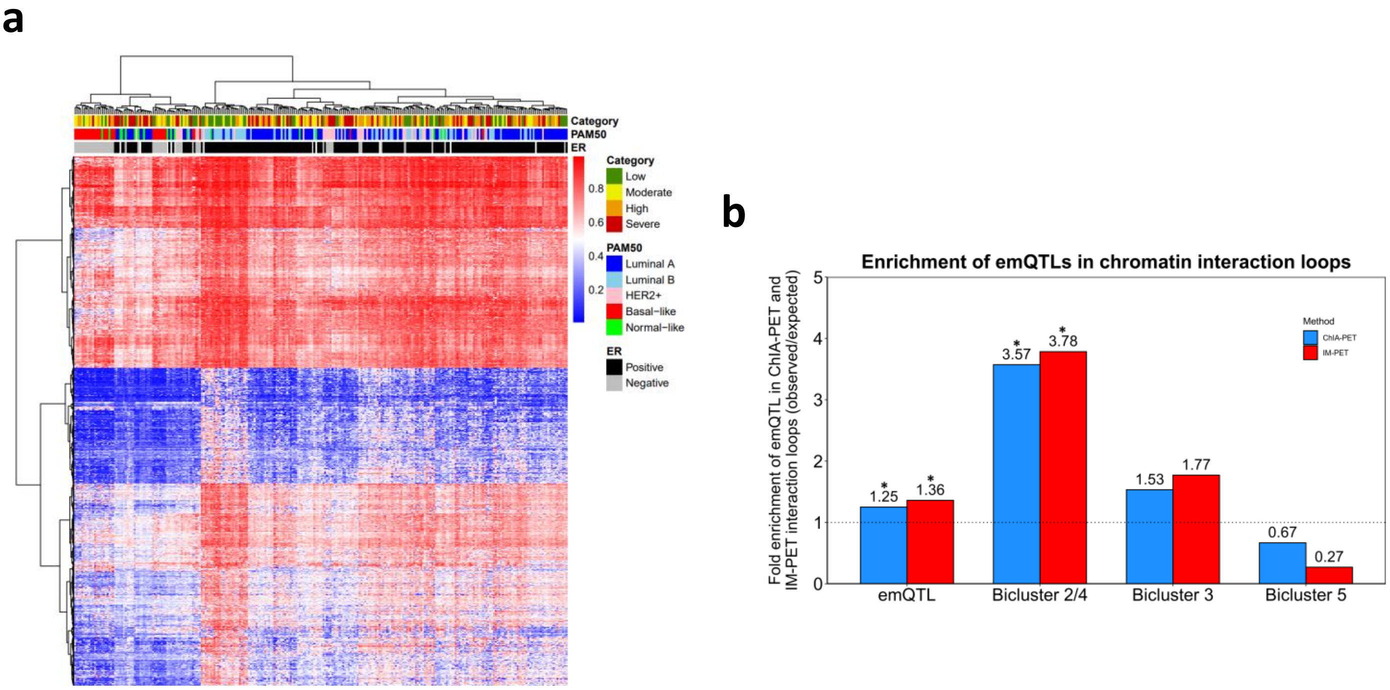


**Supplementary Figure 7. The EMT bicluster is linked to fibroblast infiltration.** Unsupervised hierarchical clustering of DNA methylation level of the 6910 EMT bicluster-CpGs in the (**a**) OSL2 (n=272) cohort. Rows represents genes while columns indicate tumor samples annotated with histopathological features including ER status and PAM50 subtype. (**b**) Bar plot showing the enrichment of emQTL-CpGs in ChIA-PET Pol2 loops and IM-PET loops for the ER+ MCF7 and ER- HCC1954 breast cancer cell lines, respectively. Bar height represents the enrichment level measured as the ratio between the frequency of emQTLs (CpG-gene pairs) found in the head and tail of a loop over the expected frequency if such overlaps were to occur at random. Statistically significant enrichments are marked with an asterisk (hypergeometric test, Benjamini-Hochberg corrected $p$-value < 0.05)

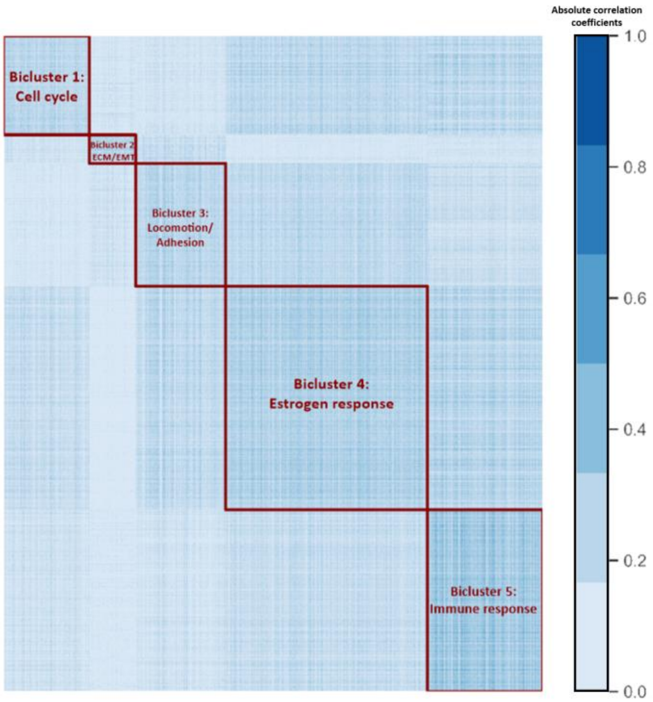


**Supplementary Figure 8. Spectral co-clustering of the absolute correlation coefficient values.** Heatmap displaying five biclusters identified by spectral co-clustering of the absolute correlation coefficient values in OSL2. Columns represents genes (*n* = 4904) and rows (*n* = 44,263) represents CpGs. Blue points indicates strong correlations between the variables while white points indicates little or no association.

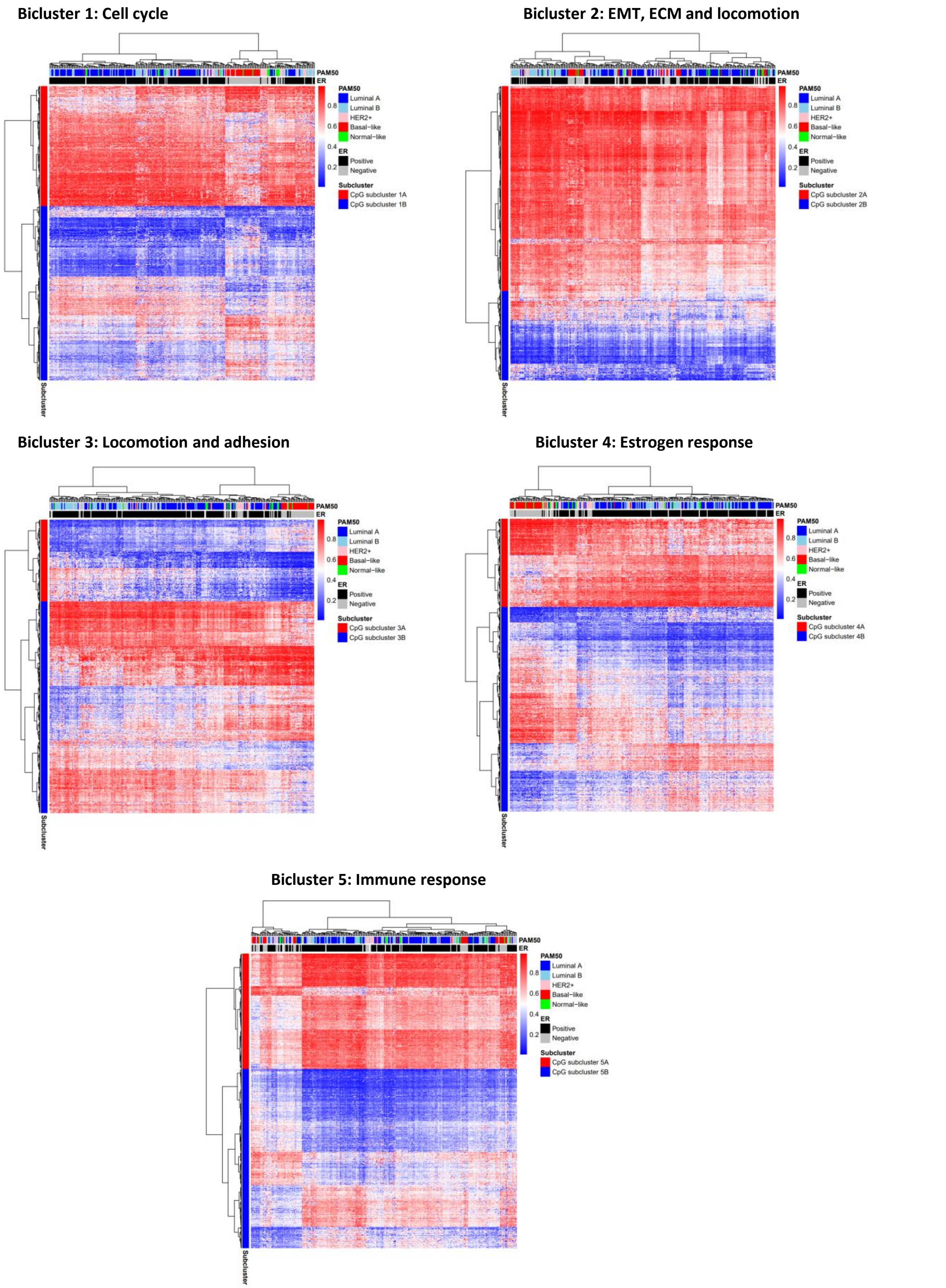


**Supplementary Figure 9.** Unsupervised hierarchical clustering of the DNA methylation levels of the CpG in each bicluster identified by biclustering of the absolute correlation coefficient values from OSL2. Rows represents CpGs while columns represents tumor samples. Histopathological features such as PAM50 subtype and ER status is shown in the annotation bar on the top. Red points indicate methylated CpGs while blue points represent unmethylated CpGs. CpGs were divided into two subclusters due to their distinct methylation patterns shown in the row annotation bar.

**Bicluster 1: Cell cycle**

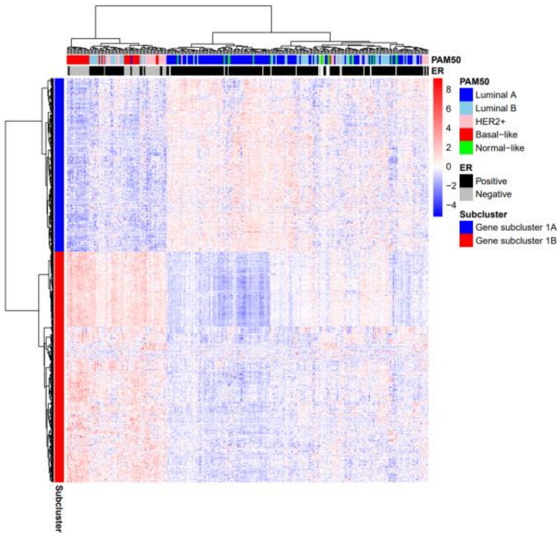


**Bicluster 2: EMT, ECM and locomotion**

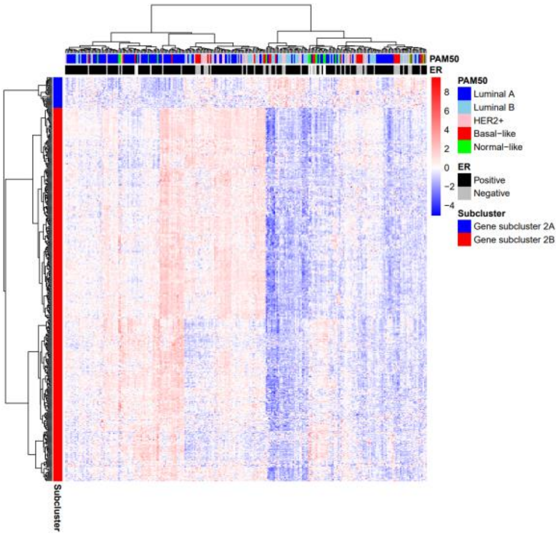


**Bicluster 3: Locomotion and adhesion**

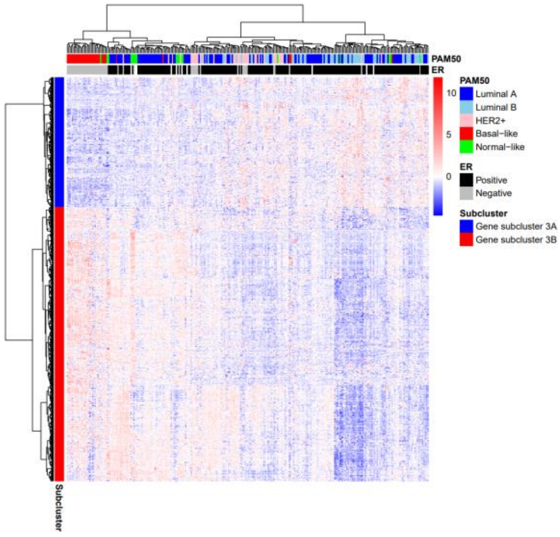


**Bicluster 4: Estrogen response**

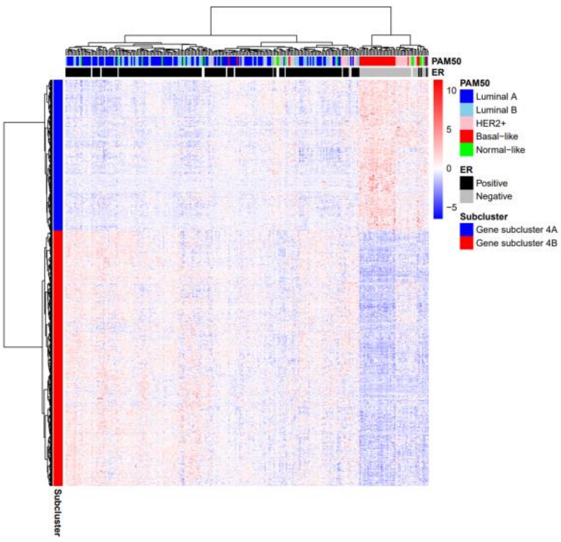


**Bicluster 5: Immune response**

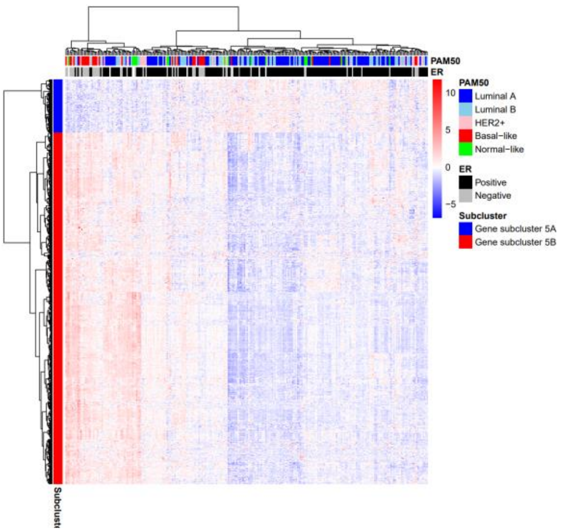


**Supplementary Figure 10.** Unsupervised hierarchical clustering of the expression levels of genes in each bicluster identified by biclustering of the absolute correlation coefficient values from OSL2. Rows represents genes while columns represents tumor samples. Histopathological features such as PAM50 subtype and ER status is shown in the annotation bar on the top. Red points indicate high expression while blue points represent low expression. Genes were divided into two subclusters due to their distinct expression patterns as shown in the row annotation bar.
